# SuperFreq: Integrated mutation detection and clonal tracking in cancer

**DOI:** 10.1101/380097

**Authors:** Christoffer Flensburg, Tobias Sargeant, Alicia Oshlack, Ian Majewski

## Abstract

Analysing multiple cancer samples from an individual patient can provide insight into the way the disease evolves. Monitoring the expansion and contraction of distinct clones helps to reveal the mutations that initiate the disease and those that drive progression. Existing approaches for clonal tracking from sequencing data typically require the user to combine multiple tools that are not purpose-built for this task. Furthermore, most methods require a matched normal (non-tumour) sample, which limits the scope of application. We developed SuperFreq, a cancer exome sequencing analysis pipeline that integrates identification of somatic single nucleotide variants (SNVs) and copy number alterations (CNAs) and clonal tracking for both. SuperFreq does not require a matched normal and instead relies on unrelated controls. When analysing multiple samples from a single patient, SuperFreq cross checks variant calls to improve clonal tracking, which helps to separate somatic from germline variants, and to resolve overlapping CNA calls. To demonstrate our software we analysed 304 cancer-normal exome samples across 33 cancer types in The Cancer Genome Atlas (TCGA) and evaluated the quality of the SNV and CNA calls. We simulated clonal evolution through *in silico* mixing of cancer and normal samples in known proportion. We found that SuperFreq identified 93% of clones with a cellular fraction of at least 50% and mutations were assigned to the correct clone with high recall and precision. In addition, SuperFreq maintained a similar level of performance for most aspects of the analysis when run without a matched normal. SuperFreq is highly versatile and can be applied in many different experimental settings for the analysis of exomes and other capture libraries. We demonstrate an application of SuperFreq to leukaemia patients with diagnosis and relapse samples.

SuperFreq is implemented in R and available on github at https://github.com/ChristofferFlensburg/SuperFreq.

## Introduction

Tracking clonal evolution within a cancer can reveal a wealth of information. In a clinical setting it can help detect the cause of relapse or drug resistance, identify early driver mutations, or track the course of metastasis[1-5]. Tracking mutations across multiple samples can also be highly informative in a research setting, including animal models of cancer, xenografts and cell lines, which often involves comparing samples over time, or across experimental conditions.

A typical analysis of multiple cancer samples from the same individual involves calling and annotating somatic single nucleotide variants (SNVs) (using methods such as multiSNV[6], VarScan 2[7], MuTect[8], SomaticSniper[9] and Strelka[10]) and copy number alterations (CNAs) (using methods such as Sequenza[11], PureCN[12] and ABSOLUTE[13]), then combining the calls within a dedicated clonal tracker (using methods such as PhyloWGS[14], SciClone[15] and PyClone[16]). The analysis will cluster mutations and produce a phylogeny, which reflects the relationship between different clones in the cancer. This multi-step process works well for an experienced user, but is sensitive to data quality issues and parameter choices. In addition, somatic SNV and CNA callers are not optimized for downstream use in clonal tracking, which makes the process of preparing the input for the clonal tracking challenging. While there are software packages that perform parts of this analysis, there is currently no integrated software that covers the entire process.

In order to address this challenge we have developed SuperFreq, a software that identifies somatic SNVs and CNAs, annotates and prioritises variants, and performs clonal tracking. SuperFreq can be used to analyse individual samples or matched samples taken over a treatment course, or from multiple sites, and is not reliant on a matched normal. We demonstrate the performance of SuperFreq by comparing it to specialised state of the art software for each step of the pipeline. We developed a complex simulation in which SuperFreq was used to perform clonal tracking on samples designed to mimic a multi-sample analysis. Finally we provide a case study in which SuperFreq was used for multi-sample clonal tracking using exome data from a patient with acute myeloid leukaemia (AML).

## Results

### SuperFreq overview

SuperFreq is an integrated analysis pipeline for cancer exomes that calls and tracks somatic point mutations and copy number alterations to reconstruct the clonal architecture of the disease. The SuperFreq workflow is presented in **Figure 1**, the input files are BAM files for test samples and reference normals, together with liberal variant calls for test samples. The outputs are annotated variant calls, including rare germline and somatic SNVs, absolute and allele aware CNAs, and clonal tracking. A major strength of SuperFreq is that it integrates many analysis components – mutation calling, quality assessment, variant annotation and clonal tracking – that are normally run separately. Combining these analytical approaches within one pipeline has major benefits in terms of the ease-of-use, but it also helps to improve the analysis, for example by allowing consistent handling of error estimates and cross checking mutation calls between samples. SuperFreq is unique in its integrated approach and as such it is impossible to benchmark with other software **(Table 1)**. Instead we provide a detailed assessment of individual components within the pipeline, such as the ability to detect somatic SNVs and CNAs, and to reconstructing clonal architecture.

**Table 1:** Properties of other mutation callers and clonal trackers in comparison to SuperFreq. identify SSNV: does the method identify somatic SNVs. SSNV w/o normal: does the method identify somatic SNVs without a matched normal. call het SNPs: does the method identify heterozygous germline SNPs for CNA calling. call CNA: does the method call somatic CNAs. CNA w/o normal: does the method call CNAs without a matched normal. subclonal CNA: can the method identify CNAs of multiple different clonalities. call clones: does the method identify clones. multisample: does the method track clones across multiple samples. track SCNA: does the method track CNAs across samples. allele aware CNA: is the method aware of different alleles affected by CNAs across samples.

| method | call<br>SSNV | SSNV<br>w/o<br>normal | call<br>het<br>SNPs | call<br>SCNA | CNA<br>w/o<br>normal | SCNA<br>clones | call<br>clones | multi-sample | track<br>SCNA |
| --- | --- | --- | --- | --- | --- | --- | --- | --- | --- |
| superFreq | yes | yes | yes | yes | yes | yes | yes | yes | yes |
| mutect2 | yes | yes | - | - | - | - | - | - | - |
| somaticSniper | yes | - | - | - | - | - | - | - | - |
| strelka | yes | - | - | - | - | - | - | - | - |
| varScan2 | yes | - | - | - | - | - | - | - | - |
| sequenza | yes | - | yes | yes | yes | - | - | - | - |
| absolute | - | - | - | yes | - | yes | yes | - | - |
| pureCN | yes | yes | yes | yes | yes | - | - | - | - |
| phyloWGS | - | - | - | - | - | - | yes | yes | yes |
| sciClone | - | - | - | - | - | - | yes | yes | - |
| pyClone | - | - | - | - | - | - | yes | yes | - |

**Figure 1:**
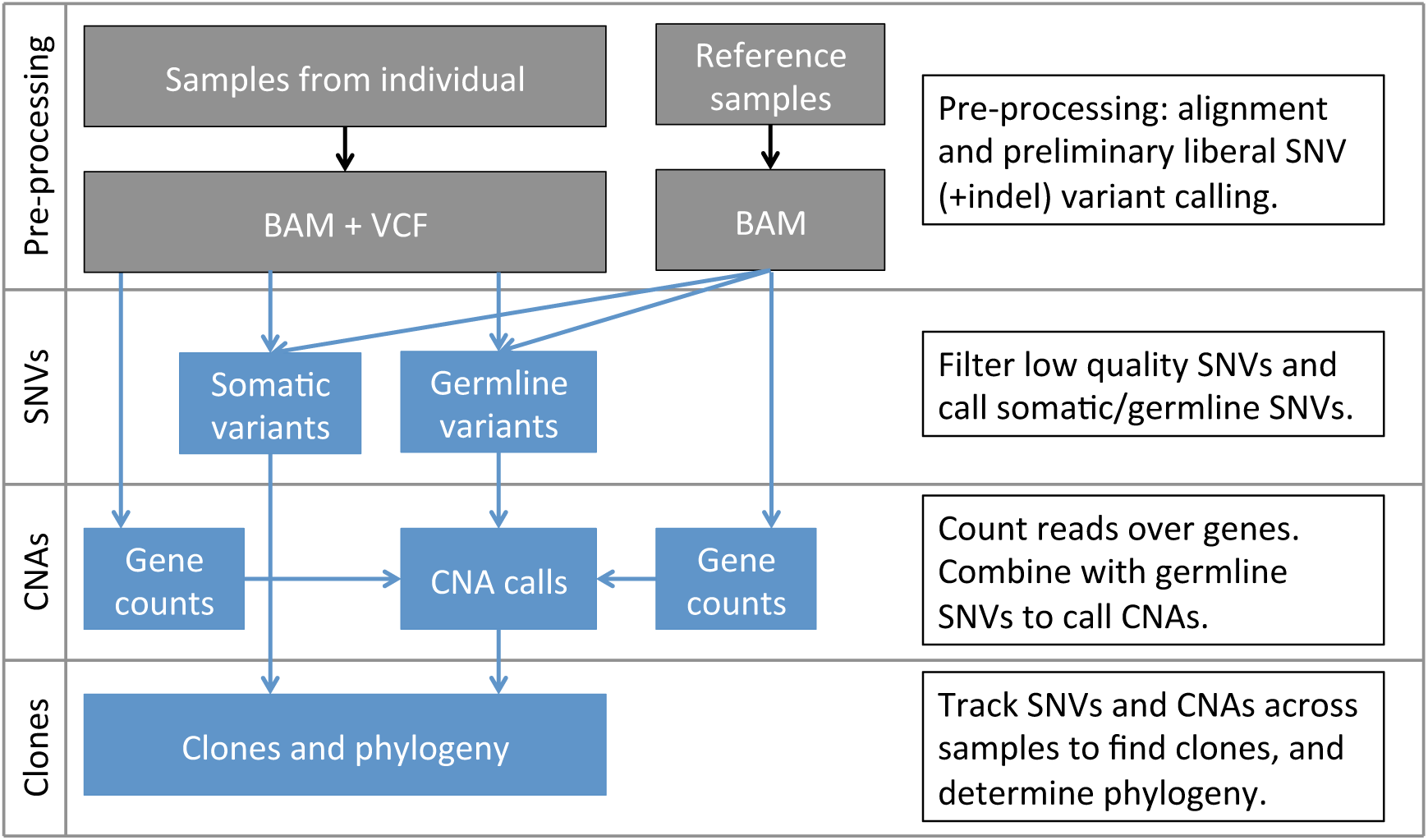
The workflow of SuperFreq. The input is aligned *BAM* files from the samples under study, and at least 2 reference normals (5-10 recommended, see methods), as well as liberal variant calls. SuperFreq filters the preliminary SNVs for artefacts using quality scores in the *BAM* file, and through comparison to the reference normals. Somatic SNVs are called from the remaining variants, while heterozygous germline SNPs are identified for CNA calling. CNAs are identified based on differences in coverage and detecting shifts in allele frequency at heterozygous germline SNPs. Finally, somatic SNVs and CNAs are analysed across samples to designate and track clones.

### Test datasets and simulation

To provide a comprehensive sample set, we randomly selected 10 cancer-normal pairs of exomes from each of the 33 cancer types included in The Cancer Genome Atlas (TCGA) and downloaded the hg38 BAM files and somatic SNV calls in VCF format from the Genomics Data Commons (GDC). A total of 26 files failed to download, because they were later excluded from the TCGA analysis, so we also excluded these cases, restricting our assessment to 304 donors. For each exome capture platform 10 reference normals were also selected, with a total of 60 reference normals used across all 33 cancer types. Details of the samples can be found in supplementary materials. We acquired CNA calls from ASCAT[17] on the matched SNP-arrays, ABSOLUTE[13] ploidies and purities from exomes and performed CNA calling on the exomes using Sequenza[11]. Simulations were performed to assess the influence of tumour purity and to generate a truth dataset for clonal tracking. To do this we performed *in silico* dilution and slicing, where reads from a cancer were substituted with those from its matched normal, either across the entire genome (dilution) or in specific intervals (slicing). This dataset provided a challenging test for clonal tracking, but is more representative of an analysis a user might face.

### Somatic SNVs

We compared the somatic SNVs identified by SuperFreq on cancer-normal pairs to calls available through the Genomics Data Commons generated with MuSE[18], SomaticSniper[9], MuTect2[8] and Varscan2[7]. In particular, we aim to minimise our false positive calls, as they can severely impact the downstream clonal tracking. First, the calls from each variant caller were compared to the consensus calls from the other four methods. SuperFreq detected a median of 91% of variants that were called by the other four callers across the 304 randomly selected donors from TCGA (**Figure 2a**). MuTect2 had a similar median (92%), while the other callers were more sensitive, with 95% for SomaticSniper, 98% for MuSE and 100% for Varscan2. Comparing to a consensus of 3 or more of the other 4 methods confirms that SuperFreq (75% recalled) and MuTect2 (79% recalled) are the two most conservative methods, while the remaining three methods recalled 90% or more of the consensus variants. However, SuperFreq only called a median of 1 somatic SNV that was not called by any other method, which was considerably lower than all other methods (**Figure 2b**). MuSE called a median of 3.5 unique variants, MuTect2 called a median of 21, while Varscan2 and SomaticSniper called 230 and 7100 unique variants respectively. Further distributions for the number of somatic variants called by permutations of two or three callers are shown in **Supplementary Figure 1** using UpSetR[19]. When looking for potential driver mutations, callers need to be sensitive to all somatic variants. However, in the context of clonal tracking false positive calls can hinder the analysis. These results therefore validate the design of SuperFreq for prioritising variants calls with low false positive rate compared with sensitivity.

**Figure 2:**
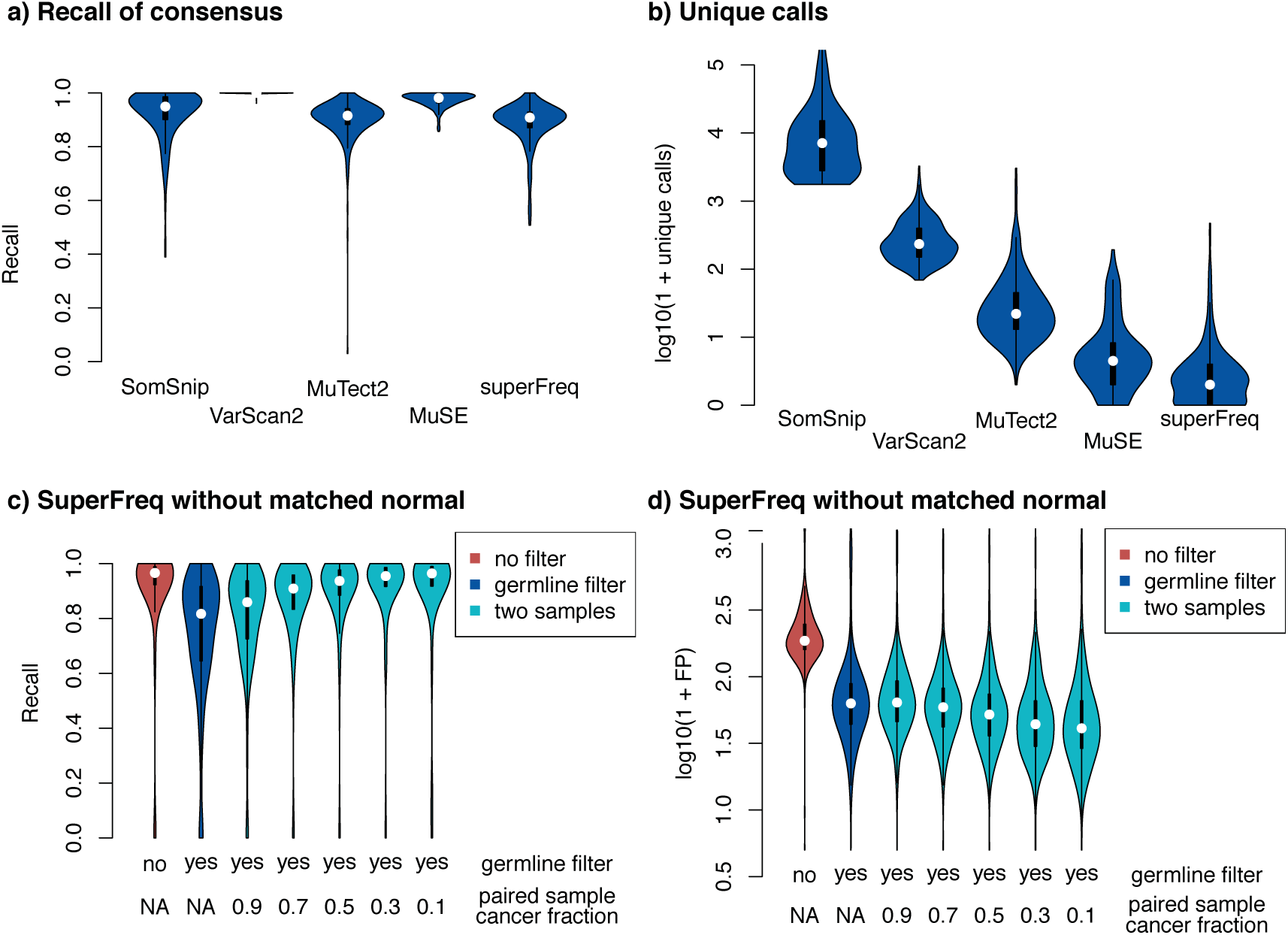
Precision and recall of somatic SNV calling across 304 TCGA participants and 33 cancer types. **a)** Recall of somatic SNVs called by the other four callers. **b)** Number of unique somatic calls generated by each caller. **c)** Recall of coding somatic SNVs from SuperFreq without a matched normal, using SuperFreq cancer-normal analysis as truth. Violins from left: cancer sample alone without filtering on the germlineLike flag, cancer sample alone filtered on the germlineLike flag, cancer sample paired with an in-silico dilution of the cancer and matched normal between 10% and 90%, filtered on the germlineLike flag. **d)** Number of false coding SNV calls in the same sample configurations.

### Somatic SNVs without a matched normal

When a matched normal is not available, SuperFreq uses population frequencies and clonal tracking to separate somatic and germline SNVs. We estimated the recall and false positive rate for calls generated without a matched normal by comparing to somatic variants identified by SuperFreq with matched normal controls. Most use cases focus on coding variants, so we restricted the benchmarking to coding variants for a more relevant assessment. When run with no matched normal SuperFreq identified a median of 97% of protein changing somatic SNVs detected in the truth set with matched normal (**Figure 2c**). However, a median of 185 additional protein changing somatic SNVs were also called (**Figure 2d**), which were largely composed of rare germline variants. We next filtered the calls using the SuperFreq *germlineLike* flag, which identifies variants that are present clonally in all samples from an individual. The germline filter reduced the median number of false calls to 62, but also lowered the median sensitivity to 82%. This drop in sensitivity is due to clonal somatic mutations being mistaken for germline variants in high purity cancer samples.

The integrated nature of SuperFreq allows it to fully utilise the information in all matched cancer samples. If multiple cancer samples are available that differ in tumour purity, they contribute together in the clonal tracking to distinguish germline variants from somatic variants. To simulate this process, we diluted the cancer sample *in silico* with sequence data from the matched normal to produce samples with lower tumour purity (10%-90% of the original cancer sample). We then analysed the original cancer sample together with the diluted sample. Adding a matched sample with 70% of the original purity and filtering on the *germlineLike* flag brought the median recall rate up to 91% with a median of 58 false calls. This shows how SuperFreq utilises even a moderate normal contamination in any of the matched cancer samples to separate somatic from rare germline variants.

### CNAs

SuperFreq monitors B-allele frequency and shifts in coverage compared to the reference normals. Segments are defined with hierarchical clustering and the clonality (cellular fraction of the sample) of each CNA is determined separately. CNAs are cross checked between all samples from the sample individual, providing a clonality estimate for each sample that can be used in clonal tracking. When cross-checking CNA calls, SuperFreq compares the direction of the signal in B-allele frequency, and splits up CNAs that affect different alleles. This allele aware CNA calling separates, for example, an AAB genotype from an ABB genotype and can help to reveal recurring events over driving genes (see Supplementary Methods, section 1.6.3).

We compared the CNA calls from SuperFreq to calls from matched SNP arrays done with ASCAT. The ASCAT calls were lifted over from hg19 with *segment_liftover[20]*. First we measured general agreement between the methods by comparing LFCs and BAFs (**Figure 3a**). Across the 292 cases where ASCAT data was available, we saw a bimodal distribution where most samples were highly concordant, but 30% of samples had agreement across less than 20% of the genome. This marked discrepancy results from different ploidy estimates. Restricting the comparison to samples with consistent ploidy estimates resulted in agreement across 95% (median) of the genome over 177 cases (blue violin in **Figure 3a**). We also used Sequenza to call CNAs from the exomes and repeated the comparison to the ASCAT calls (**Figure 3a**), and found similar results. Sequenza has a consistent ploidy with ASCAT in more cases (198), but had lower overall agreement with ASCAT across the genome (90% median) for these cases. SuperFreq relies on the reference normal samples for the LFC estimates, and only uses the matched normal to identify heterozygous germline SNPs, so we did not expect to see major differences in performance with or without a matched normal set. Indeed, when we performed the SuperFreq analysis without matched normals we found a maintained agreement with ASCAT across 95% (median) of the genome in 181 samples with concordant ploidies (**Figure 3a**).

**Figure 3:**
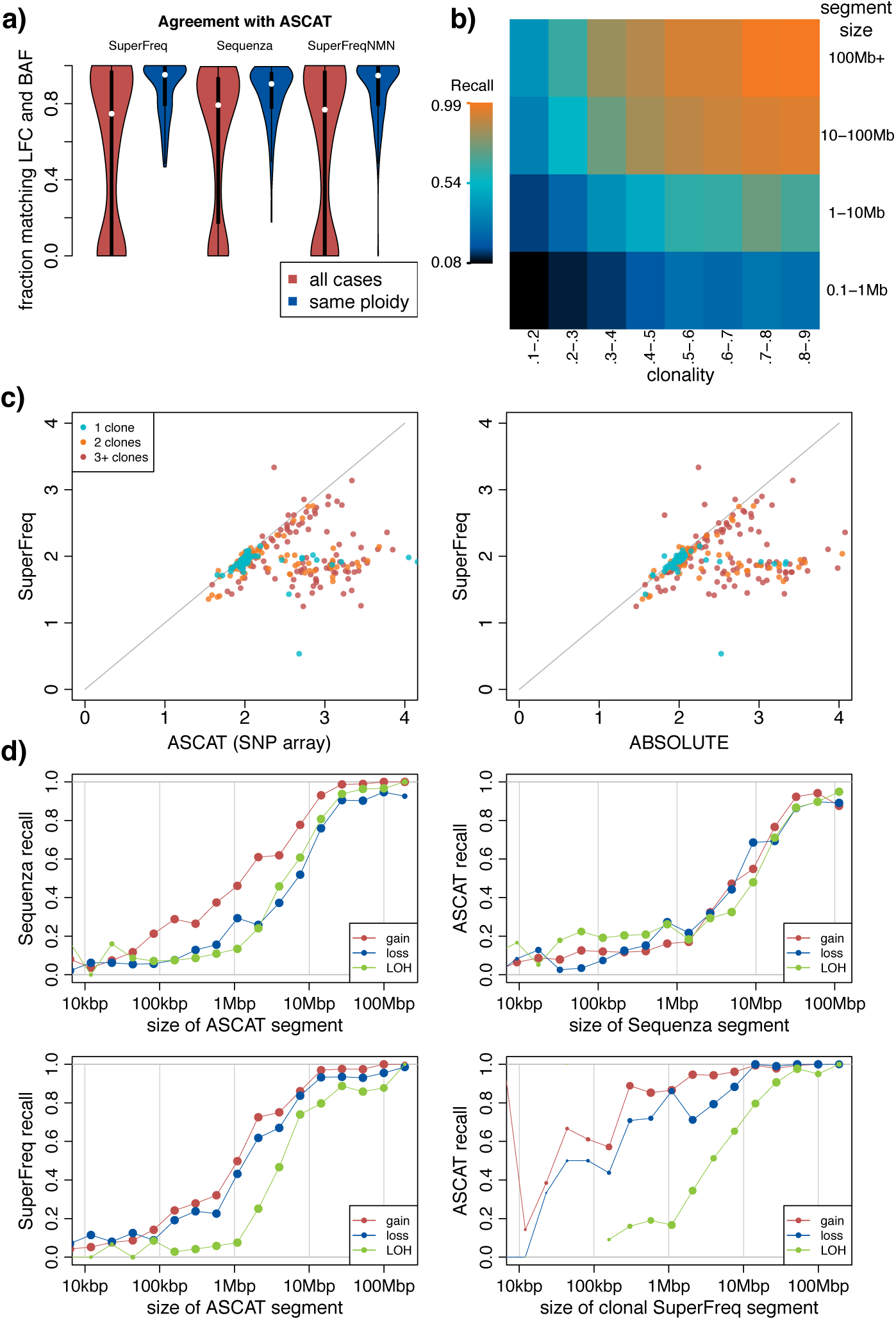
Comparison of somatic CNA calls for 304 TCGA participants across 33 cancer types. **a)** Distribution across participants of the fraction of the genome that agrees with ASCAT. Agreement is defined as LFC and BAF within 0.145 and 0.05 respectively, roughly corresponding to a 20% clonality gain or loss. The blue violins show only participants with a ploidy estimate within 0.2 of ASCAT. **b)** SuperFreq recall of somatic CNAs in diluted and sliced samples analysed without the matched normal, with the original cancer-normal analysis used as truth. Each bin is based on at least 100 copy number segments. **c)** Comparison of ploidy estimates between ASCAT, ABSOLUTE and SuperFreq, coloured by number of somatic clones called by SuperFreq. Samples were excluded if no somatic clone was called. **d)** Recall of gain, loss and CNN-LOH binned by size of the segment, limited to participants where the ploidy agrees within 0.2 between the methods.

**Figure 4:**
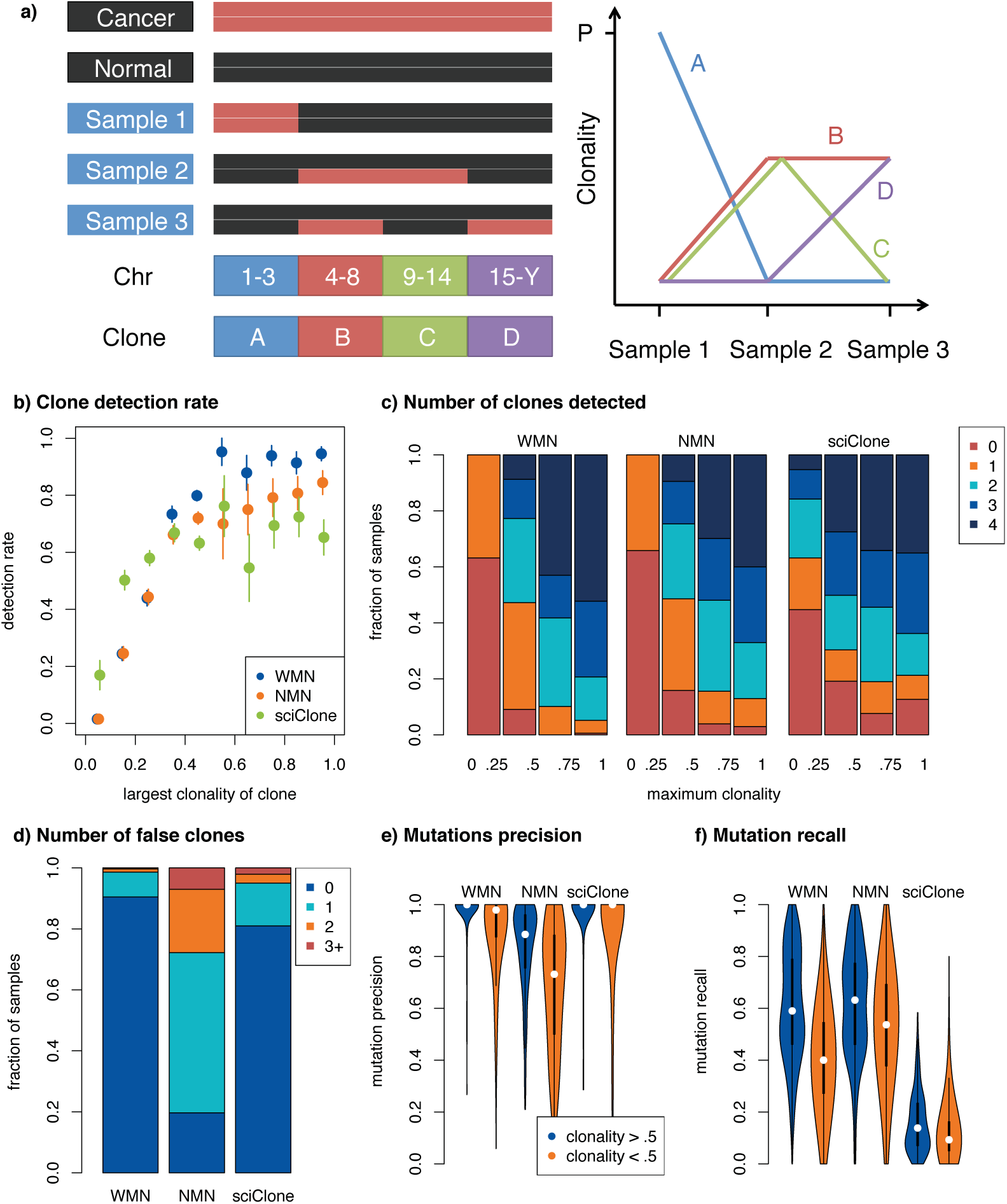
Precision and recall of SuperFreq clonal tracking. **a)** Overview of simulations. As illustrated on the left, the genome is divided into four regions (chr 1-3, 4-8, 9-14 and 15-Y), and the cancer and normal samples are blended to create three samples that contain four clones supported by mutations that reside in different regions of the genome. The expected clonalities across the three samples are shown to the right, where P denotes the purity of the cancer sample. **b)** Sensitivity to find clones with a matched normal (WMN) or no matched normal (NMN) in SuperFreq and SciClone, as function of maximum clonality. **c)** Recall of the four simulated clones, binned on the purity of the original cancer sample. **d)** Number of false clones called. **e)** Fraction of mutations associated with a clone originating from the expected chromosomes in panel **a. f)** Fraction of mutations called in the cancer-normal recalled by the called clones.

We investigated the differences in ploidy estimates between methods. SuperFreq does not explicitly call a purity, so for comparison purposes we calculate ploidy as the average number of haploid genomes across all cells in the sample, excluding sex chromosomes. While SuperFreq and ASCAT agree on the majority of the cases, there are 60 samples where ASCAT calls a significantly higher ploidy (>1 larger) (**Figure 3c**). It is difficult to accurately estimate ploidy using SNP arrays or exomes and in the absence of orthogonal measurements we cannot determine which method is most accurate. It could be that the SuperFreq estimates are overly conservative, or that ASCAT is adjusting ploidy to accommodate subclonal CNAs. Most (90%) of the cases with discrepant ploidy estimates had subclones in the SuperFreq analysis. Comparing SuperFreq to Sequenza revealed a similar pattern (**Supplementary Figure 2-4)**. TCGA cases typically do not have orthogonal ploidy information, but the TCGA-AML cohort has matching cytogenetics which provides a more reliable ploidy measurement. AML has a very low rate of CNAs, and none of the 194 cases with cytogenetic data has more than 53 chromosomes. Nonetheless, ASCAT calls a ploidy >3 in 11 cases (6%), suggesting some high ploidy calls are artefactual. ABSOLUTE is another method that calls absolute copy numbers that can account for subclones. When ABSOLUTE was applied to the AML dataset it found a ploidy >3 in only 2 out of 118 AML samples (2%). We next compared the SuperFreq ploidy estimates to those from ABSOLUTE for the TCGA test cohort, which were available for 304 samples (**Figure 3d**). We found that there was a modest improvement in the agreement, but there were still 49 samples (16%, 37 samples overlap with ASCAT) where ABSOLUTE called a ploidy at least 1 larger than SuperFreq. A better truth data set is needed for comparison of ploidy from CNA callers.

We next assessed the resolution of the CNA calls from SuperFreq by looking at the sensitivity based on the size of the event. For this purpose, we classified each segment into gain (copy number larger than 2), loss (copy number smaller than 2) and CNN-LOH (copy number neutral loss of heterozygosity) based on the calls from ASCAT, SuperFreq and Sequenza. We performed a pair-wise comparison between methods, where we used one method as truth and measured the fraction of the truth segment that was covered by segments with the same class of copy number event in the second method. We first compared the more established methods, ASCAT and Sequenza, binning by the size of the truth segments (**Figure 3d**). We found that CNA calls of all three classes had reliable recall when the size of the truth segment was >10 Mb. For CNAs smaller than 10 Mb recall dropped rapidly; below 1Mbp both Sequenza and ASCAT recall very few of the CNAs of the other method. SuperFreq has a similar recall of ASCAT segments as Sequenza (**Figure 3d**), also with low recall of small CNAs. SuperFreq calls significantly fewer small segments than Sequenza (median 1 segment below 1Mbp per sample, compared to median 16), which results in higher accuracy for small copy number segments. Indeed, ASCAT agrees on the majority of small SuperFreq gain or loss calls down to 100kbp (**Figure 3d**). This suggests that SuperFreq has a very low rate of false calls for gain and loss events. SuperFreq calls subclonal copy numbers, so when SuperFreq acted as truth, we restrict it to clonal CNAs, with a clonality > 0.5, and found good agreement for events greater than 0.5 Mb (**Figure 3d**). Without restrictions on the SuperFreq truth, we found decreased recall for large segments (>10Mbp) (**Supplementary Figure 5**). This may seem counterintuitive but is expected when considering that ASCAT does not call subclonal events, and that larger segments provide more power to identify subclonal CNAs. Indeed, ASCAT recalls close to 100% of SuperFreq’s large CNA calls with clonality above 0.5. Comparing SuperFreq to Sequenza shows similar behaviour (**Supplementary Figure 5**), with the difference being that Sequenza has higher recall of small CNN-LOH. This may indicate that these small CNN-LOH calls are true calls that are missed by the SNP array, or it could be related to exome-specific artefacts. A comparison between SuperFreq with and without a matched normal shows high recall for loss and gain events across all resolutions, but with worse recall for CNN-LOH, which relies on accurate determination of germline SNPs.

**Figure 5:**
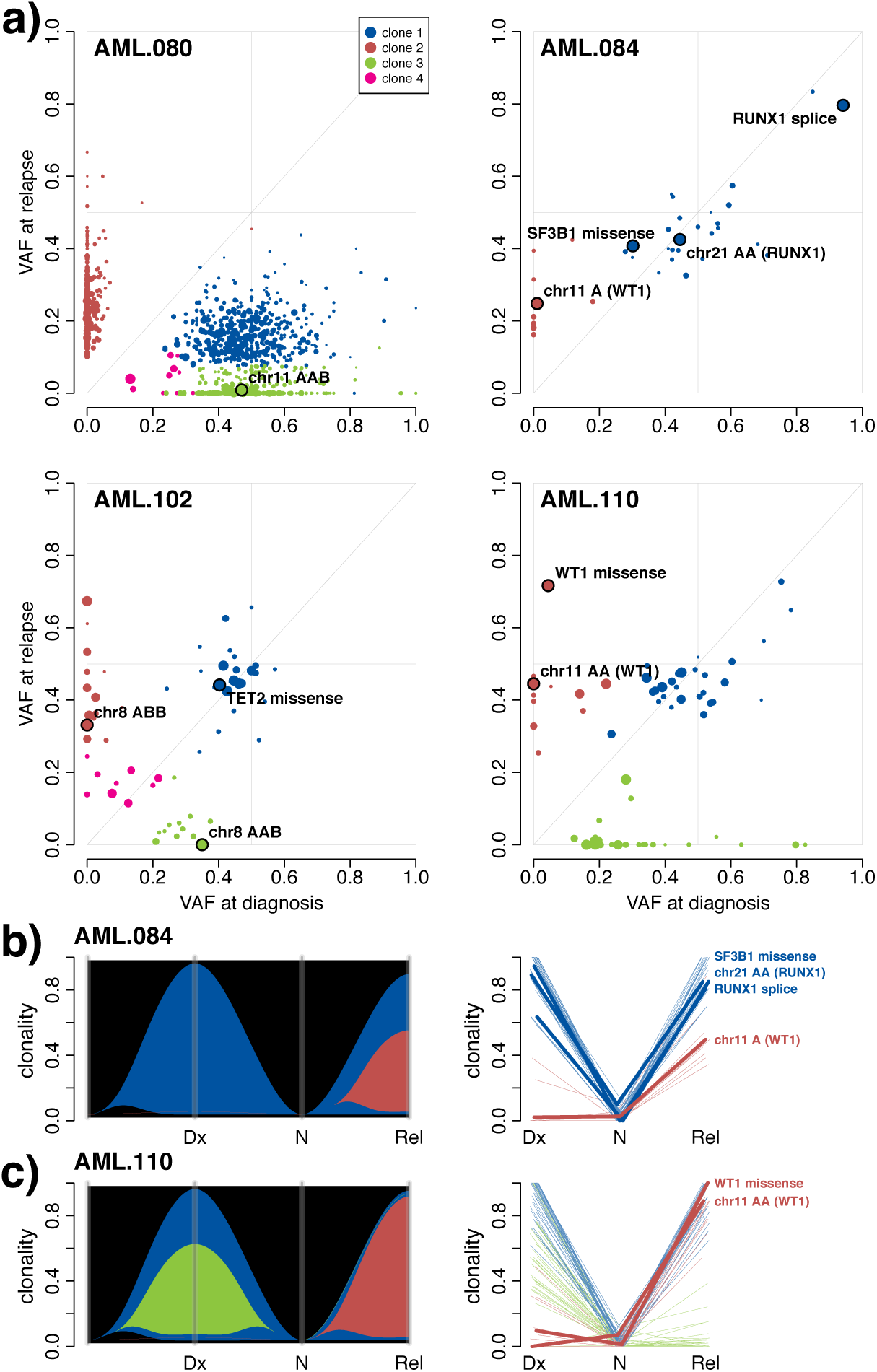
Assessment of clonal structure in AML using SuperFreq. **a)** Clone scatter plots of four AML patients. The VAF is shown for all somatic point mutations that are tracked by SuperFreq, coloured by the clone they are assigned to. Selected mutations are highlighted, including CNAs that are assigned a VAF of half of the clonality. Point size is used to indicate read depth, with larger points reflecting higher coverage. River plots and line plots showing the progression of the clones across the Diagnosis (Dx), sorted normal (N) and relapse (Rel) samples for **b)** AML.084 and **c)** AML.110, key mutations are highlighted in the line plots.

In order to assess SuperFreq’s sensitivity to CNAs covering small genomic regions and those present at low purity, we diluted the cancer sample with the matched normal to simulate lower purity CNAs. We also generated sliced samples in which reads from set regions of the cancer sample replaced those in the normal sample. In this way we created samples with CNAs spanning specific genomic regions, where we could control the size, and could also approximate lower tumour purity. Using the SuperFreq cancer-normal calls as truth, we measured the rate of recall from SuperFreq run on sliced and diluted samples as function of the size and clonality of the CNA. When using the matched normal to dilute or slice the sample, we cannot use the same matched normal for analysis without introducing bias. For this reason we restricted this assessment to cancer-only analysis, calling CNAs from the mixed sample without a matched normal, which avoids this bias. We found that above 10Mbp and 30% clonality, almost all CNAs are called, and there is then decreasing sensitivity for smaller events and lower purity. (Figure 3d). The dilution series also provide a truth for the clonality of the copy numbers, which are expected to be proportional to the cancer fraction (see example in Supplementary Figure 6).

**Figure 6:**
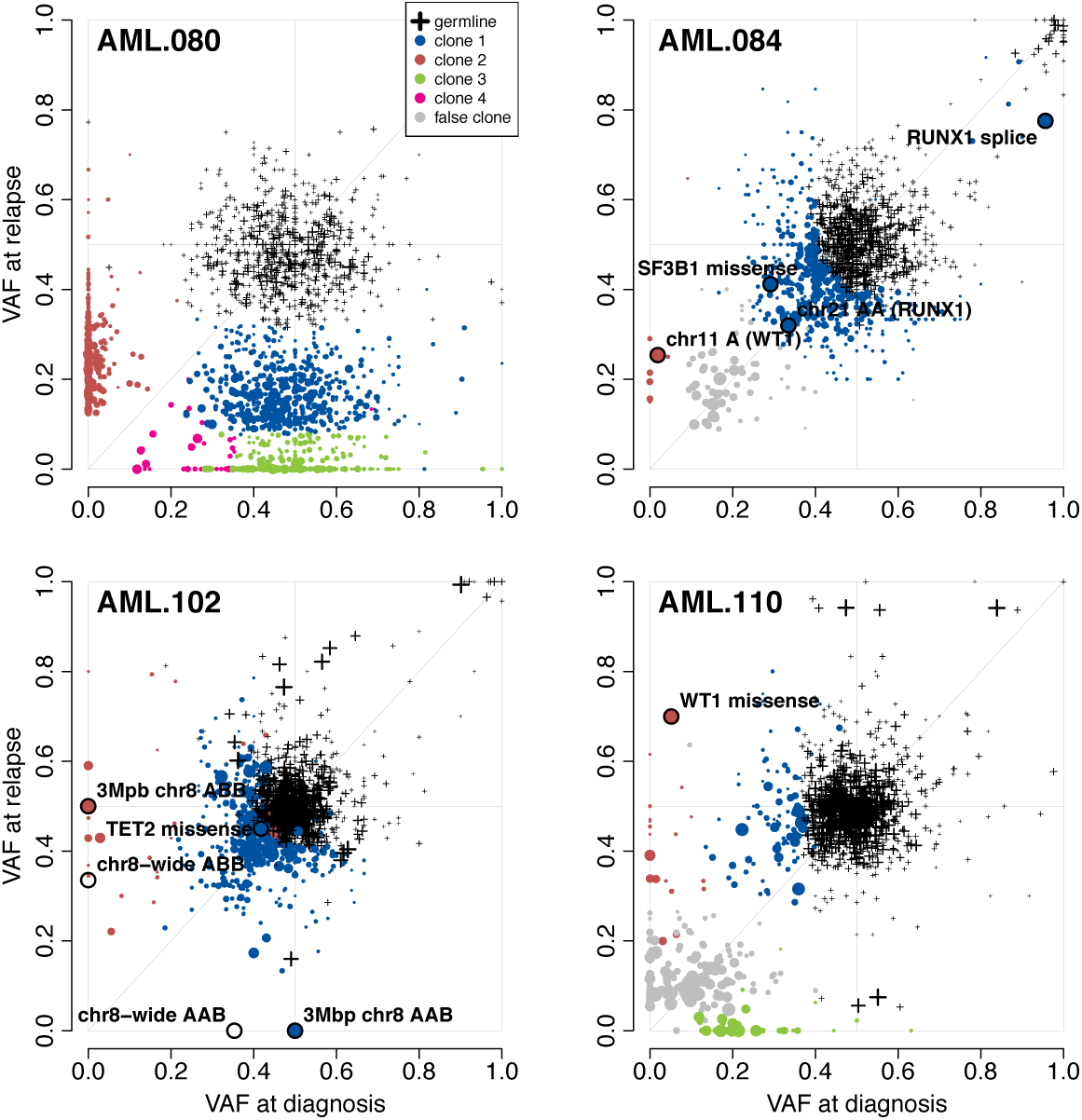
Assessment of clonal structure in AML using SuperFreq without matched normal controls. Clone scatter plots of four AML patients, as in Figure 5, generated without using the matched normal sample. We include the germline clone to illustrate how this clone absorbs germline variants, as well as some somatic variants that have high clonality in both samples. While this is done successfully in AML.080 and AML.110, the two other samples have a large number of variants called as a somatic clone that are not identified in the cancer-normal analysis. AML.084 and AML.110 identify clones at low clonality in both samples that are not seen in the cancer-normal analysis, shown in grey.

### Clonal tracking

Existing clonal trackers are generally run on highly curated sets of somatic SNVs and CNAs. SuperFreq, by contrast, works with input variant calls made with permissive settings and performs an internal quality assessment. This quality assessment relies on variant characteristics derived from the BAM file. To generate a test dataset for clonal tracking with SuperFreq, we created three matched cancer samples for each TCGA participant by diluting (replacing cancer reads with normal reads) and slicing (replacement of cancer regions with those from the normal) the BAM files from the cancer (**Figure 4a**, see methods). Dilution and splicing was used to create four distinct clones in each sample set, that carry mutations in different genomic regions, which we could use to estimate recall and false positive rate. Only participants that had at least one cancer clone detected in the cancer-normal analysis were included (289 out of 304), but there were otherwise no restrictions on the number of mutations split across the four clones. The analysis was performed both with and without the matched normal control.

We analysed the matched samples with SuperFreq and measured the rate of recall of clones, as a function of the maximum clonality (**Figure 4b** and **c**), where clonality is the cellular fraction of the sample. We also assessed how many mutations were correctly attributed to each clone, based on the chromosome harbouring the mutation. An example participant is shown in **Supplementary Figures 7-9**. As we only have access to one normal and one cancer sample, we cannot avoid biases between samples, where the same reads will represent the cancer or normal cell fractions in all samples. Across 289 test datasets generated from TCGA samples, we found that SuperFreq detected 93% of the simulated clones with a maximum clonality above 50% (**Figure 4b**). When considering participants with tumour purity above 75%, SuperFreq detected all four clones in 52% of cases, and three or more clones in 79% of cases, (**Figure 4c**). SuperFreq had a lower false positive rate, calling a false clone in less than 10% of cases (**Figure 4d**). When considering clones that were correctly identified, we can measure precision and recall of the mutations contributing to the clone. For clones above 50% clonality, SuperFreq recalled a median of 59% of the mutations with median 100% precision (**Figure 4e-f**). We next assessed clonality calling in the absence of a matched normal control. In this analysis SuperFreq had lower recall for clones above 50% clonality, dropping from 93% to 80%. When considering cases with high tumour purity, SuperFreq still recalled three or more clones in 67% of cases. There was a marked increase in the fraction of cases in which a false clone was called, increasing from 10% to 80%. The median recall rate of mutations remained similar, but with a drop in median precision from 100% to 89%.

For comparison of the clonal tracking step, the CNAs and somatic SNVs called with a matched normal were also analysed by SciClone: a dedicated clonal tracker. SciClone has relatively strict requirements for somatic SNVs, with a default requirement for at least 10 high quality SNVs in regions with normal diploid copy number. As some of the cases did not meet this requirement, the default filters were gradually relaxed until the algorithm could be executed. Note that this does not constitute a comparison of the entire SuperFreq pipeline, only the last clonal tracking step. Performance depended strongly on the SciClone settings and details of the relaxed requirements. With the best analysis scheme, SciClone achieved a lower sensitivity of 67% at clonalities above 50% (**Figure 4b**), but had higher sensitivity at low clonality. Similarly for the number of clones detected, SciClone detected fewer clones at high clonality (detecting all 4 clones in 35% of cases above 75% clonality), but displayed higher sensitivity at low clonality. SciClone showed a higher false positive rate, calling a false clone in 19% of cases with matched normal.

### Application: clonal tracking in AML

To demonstrate how SuperFreq can be applied for clonal tracking, we present an analysis of a cohort of patients with relapsed AML [1]. For these patients, samples were available at diagnosis, at relapse and from purified lymphocytes, which could be used as a matched normal control. We ran SuperFreq using default parameters with up to 10 normal samples from the same centre as reference normals. We found that SuperFreq could identify somatic SNVs and CNAs and reliably distinguish diagnosis specific events, relapse specific events and those present at both time-points (**Figure 5**). Overall the clonal structure predicted for each sample was consistent with the published results generated with sciClone[1], with the exception of low abundance clones that were more variable. These results were in keeping with our TCGA simulations (**Figure 4**). There were some cases with marked differences in predicted clonal structure, for example SuperFreq identified a diagnosis-specific clone in AML.102 that was not defined in the original analysis. This diagnosis-specific clone is supported by a high confidence CNA call, a chromosome wide gain on chromosome 8 (designated AAB in **Figure 5A**). Chromosome 8 was also gained at relapse in that patient, but allele specific tracking showed it was the alternate allele (designated ABB) (highlighted in **Supplementary Methods Figure 7)**. This example shows how allele specific tracking can help to define recurrent CNAs to aid identification of important functional mutations.

We provide a more detailed review of AML.084 and AML.102. For AML.084 a single, dominant cancer clone (blue) was detected at diagnosis (**Figure 5B**). Candidate somatic variants are prioritised, based on variant effect and comparison to the COSMIC database. In this case four mutations were detected in COSMIC census genes, including a splice site variant in *RUNX1* and a hotspot mutation in *SF3B1* (K700E). We also detected a CNN-LOH event on chr21 (designated chr21 AA), which extends over 31Mbp and includes *RUNX1*. The *RUNX1* variant has a VAF significantly larger than 50%, consistent with loss of the wildtype allele. At relapse, AML.084 exhibits a new subclone present at around 40% clonality, which carries five protein altering SNVs, together with loss of a 10Mbp segment on chr11. Closer examination of the CNA on chr11 revealed loss of the tumour suppressor gene *WT1*. When we examined AML.110 we found that it had also acquired mutations in *WT1* at relapse, both a missense mutation (D464N) and CNN-LOH. These cases highlight an important feature of SuperFreq, which is the tracking of CNAs, together with correction of VAF for local copy number, improves the accuracy of clonal tracking and aids identification of important genes.

We repeated the end-to-end SuperFreq analysis of these four cases without the matched normals (**Figure 6**). Most of the clonal structure and key mutations were recalled, but with an increase in false calls. We note that the four point mutations in AML genes are called and tracked, as well as the RUNX1 CNN-LOH. However, the CNN-LOH over WT1 was missed owing to the very high clonality (>90% of cells), which made it difficult to distinguish. In the absence of the matched normal we note the increased number of false somatic clones. These false clones result from the inclusion of low quality SNV calls, they are generally present at low clonality and tend to show similar clonality at diagnosis and relapse.

In the absence of a matched normal SuperFreq retained high recall of somatic variants (91%), with 9% of variants excluded because of their presence in population data bases. Without a matched normal we saw inflation of the total candidate somatic variant count due to inclusion of rare coding variants (average 226 coding variants per sample), but application of the germline filter reduced this by approximately 70% (average 68 coding variants per sample) (**Supplementary Table 1).** Using the *germlineLike* filter, the recall dropped to 87% total, but this assessment was skewed by inclusion of AML.080, which has a very high mutation load (**Supplementary Table 1**). The other three patients have recall below 50% with this filter, due to the high purity of the cell sorted cancer samples. This highlights an important point, that caution must be taken when analysing samples with high tumour content in the absence of a matched normal control because of the overlap in the distribution of the VAFs for somatic and germline variants (**Figure 6**). In this case the contaminating normal tissue helps to separate somatic mutations from rare germline variants.

## Discussion

Recapitulating the evolutionary history of a cancer from sequencing data can be tremendously insightful, but it is technically challenging. Selecting high quality somatic variants appropriate for clonal tracking is a significant barrier. This can be further compounded by technical imperfections in the sample data, or the absence of a high quality matched normal. SuperFreq addresses these challenges; it provides a single analysis that performs somatic SNV filtering, CNA calling and clonal tracking, without requiring a matched normal. Variants are annotated with their effect on proteins and compared to population and cancer databases to aid interpretation and to highlight potential driver mutations.

A key aspect to maximise sensitivity while limiting false calls throughout the analysis was to maintain accurate error estimates. Using a single value as the error estimate allowed us to propagate the error throughout all the steps of the analysis and allowed us to account for multiple error sources. The error estimates inform multiple steps in the analysis, such as copy number segmentation and calling, mutation clustering, and classification of anchor mutations.

To show that the workflow is robust, we analysed exomes with SuperFreq across 33 cancer types in TCGA. This ensured heterogeneity in somatic SNV and CNA mutation rate, as well as technical biases, and allowed us to assess performance of SuperFreq in a wide range of settings. We showed that SuperFreq recalled more than 90% of the somatic SNVs and CNAs (those larger than 10Mbps) identified by a consensus of other methods, and maintained a similar level of performance without a matched normal, albeit with more false positive calls. Some aspects of copy number calling remain challenging, including estimating ploidy, detecting small events and accounting for overlapping subclonal CNAs, highlighting the need for detailed orthogonal testing to validate calls. In the absence of a matched normal, we found that having matched cancer samples with different levels of purity also improved the precision and recall of somatic SNVs.

To assess the ability of SuperFreq to reconstruct a clonal history, we developed an innovative approach where we sliced and blended data from TCGA samples to produce sets of samples with an expected clonal structure. Partitioning mutations to specific genomic intervals aided the detection of false positive calls. This was expected to be a challenging data set to analyse, but SuperFreq recovered 93% of clones above 50% clonality, with fewer than 10% of cases having false clones. The frequency of false clones increased markedly in the absence of a matched normal control, which means extra caution should be taken when working without a matched normal control. While we found that SuperFreq can reliably perform a fully automated analysis of somatic SNVs and CNAs followed by clonal tracking, we recommend orthogonal testing to validate predicted clonal structures.

SuperFreq was designed to detect and track somatic mutations in exomes, and it has been applied to study breast cancer metastasis[2, 21], lung cancer xenografts[22], gastric cancer organoids[23], and myeloid leukaemia[24]. SuperFreq is highly versatile and it has since been applied to study small capture sets[25] and low pass whole genomes[26]. We want to draw attention to De Mattos-Arruda[21], where multiple pipelines, including SuperFreq, were used in parallel for clonal tracking, and the authors found overall quite consistent results between methods, providing an independent example of how SuperFreq can be applied for an end-to-end analysis on real data. Further improvements in how we benchmark clonal tracking methods is an important consideration for the field. We are currently optimising performance to allow routine analysis of whole genomes and have also produced promising results when applying SuperFreq to transcriptomes.

## Methods

Section 1 of the supplementary methods contains a comprehensive description of the SuperFreq algorithm. This paper describes version 1.0.0, the current version of the software is available at https://github.com/ChristofferFlensburg/SuperFreq.

### Initial data inputs to SuperFreq

The input to SuperFreq is a set of indexed BAM files for the samples and reference normals, together with metadata of the samples and the reference genome to which the samples were aligned. We have found that 5-10 normal samples are generally sufficient. We found performance did drop with a lower number of normal samples (**Supplementary Table 2**), but in our experience the quality of the normal samples, in terms of their ability to reproduce technical artefacts found in the test samples can be more important than using a larger number of control samples. “We expect good recall for heterozygous clonal variants at >30x read depth, but recall will drop as coverage diminishes.”

### SNV and small indel quality control

SuperFreq first performs preliminary variant calling on each sample with liberal settings using Varscan. This step can be skipped if the user provides VCF files with preliminary variant calls directly. SuperFreq shares the preliminary variants across all samples and filters variants using base quality, mapping quality, and strandedness. Variants present in the reference normals are removed from the analysis of somatic SNVs, but common population polymorphisms are retained for CNA calling. This is described further in section 1.2 of the Supplementary methods.

### Somatic SNV calling and annotation

Variants are identified as somatic if they have a significantly higher variant allele frequency (VAF) in the cancer compared to the normals. If there is a matched normal sample we perform a Fisher exact test on the number of variant and reference reads between the cancer and the matched normal. In the absence of a matched normal, a filter is applied to exclude variants with a population frequency > 0.1% (dbSNP[27] and ExAC[28]). Candidate somatic variants that cluster with the germline in the clonal tracking are marked with the *germlineLike* flag. The somatic SNV assessment is summarised in a quality score *somaticP* between 0 and 1 reflecting the confidence that the variant is somatic. For downstream analysis of somatic variants, we typically use *somaticP* > 0.5 as cut-off, but it can be adjusted to favor precision or recall. Somatic SNVs are annotated using Ensembl Variant Effect Predictor[29], and candidate driver mutations are highlighted through comparison to the Catalogue Of Somatic Mutations In Cancer[30]. The details of somatic SNV identification in SuperFreq is described in the Supplementary Methods, section 1.3.

### CNA calling

SuperFreq uses read coverage and B-allele frequencies (BAFs) at heterozygous germline variants to call CNAs. FeatureCounts[31] is used to determine the read count over each capture region (exon) for each sample. The read counts are corrected for GC-bias and correlations between total read count and LFC with respect to the reference normals (Supplementary Methods, figure 1 and 2). SuperFreq runs limma[32]-voom[33] with sample weights[34] on the bias-corrected counts to test for an increase or decrease in coverage indicating a CNA. Each sample is compared, one-against-many, to the reference normals, resulting in a log fold change (LFC) in read depth and t-statistic for each region. SuperFreq exploits the expected property that most adjacent capture regions will share the same true LFC, i.e. that the number of true copy number breakpoints is much smaller than the number of genes. With that assumption, a median difference between adjacent capture regions larger than expected from the limma-voom variance estimates is a sign of underestimated variance, which is corrected by adding a constant to the variance estimate, as shown in figure 3 of the Supplementary Methods. By using the median difference, we are not sensitive to the small fraction of neighbours that span a copy number breakpoint. The analysis of the read counts is described comprehensively in section 1.4 of the Supplementary methods.

In addition, heterozygous germline SNPs are identified for use in CNA calling and used to determine the B allele frequency (BAF). If a matched normal is present, common population variants that are observed to have close to a 50% variant allele frequency (VAF) in the matched normal are used. If no matched normal is present, then variants with > 1% population allele frequency with a sample VAF between 5% and 95% are used. Each copy number segment is tested for balanced allele frequency using a log likelihood ratio approach, as described in the supplementary methods (section 1.5 and figure 4).

Finally, the genome is segmented into copy number regions based on the coverage LFC and BAF. The capture regions for each gene are merged and hierarchical clustering is performed. The most similar adjacent segments are merged recursively, with a distance measure comparing LFC and BAF. The ploidy of the sample is then determined from the ratio of the coverage LFC with respect to the reference normals. Different segments are assigned clonality (sample cellular fraction) and copy number call independently. Each segment is assumed to only have a single copy number alteration. This process is illustrated in the maypole plot in **Supplementary Methods, Figure 6**, where ploidy corresponds to a constant shift along the x-axis, and is described in the supplementary methods section1.6.

### Clonal tracking

The clonality (sample cellular fraction) of each somatic SNV is calculated based on the VAF, accounting for local copy number. The clonality of each CNA is tracked over samples, and alterations affecting different alleles are split into separate mutations (e.g. AAB and ABB genotypes), see supplementary methods section 1.6.3. The SNVs and CNAs undergo hierarchical clustering based on the clonality and uncertainty across all samples. The resulting clusters are required to be consistent with a phylogenetic tree. Specifically we require clonal unitarity: that the immediate subclones are not allowed to have a significantly higher summed clonality than that of the parental clone. Inconsistencies are resolved by removing the clone scoring highest in a set of properties typical of false clones, such as constant clonality, high proportion of indels compared to SNVs, or few supporting mutations. The clustering is initially performed with only high confidence somatic mutations. Mutations with lower confidence are then added to the most similar cluster, or discarded if no sufficiently similar cluster is found. The details are available in section 1.7 of the supplementary methods.

### Resource usage

For a cancer-normal pair of exomes with 10 reference normal samples, SuperFreq typically runs in 3 hours on 5 cpus, using 20Gb of memory and creating 400MB of data and plots. Runtime and memory usage vary depending on multiple factors, such as number of germline variants, number of somatic mutations, library size and levels of noise in the studied samples.

## Availability

SuperFreq is available as an R package on github: https://github.com/ChristofferFlensburg/SuperFreq/ The data underlying the results presented in the study are available from the database of Genotypes and Phenotypes and the Genomic Data Commons, after approval of each respective data access committee. We have made processed data available, to enable reproduction of results and figures, however, germline variant calls have been censored. Results from the TCGA analysis and code to reproduce the figures are available at: https://gitlab.wehi.edu.au/flensburg.c/SuperFreqPaper

## Ethics Statement

This study involves the analysis of published cancer genomics data sourced from the Genomic Data Commons. Research datasets are detailed in the acknowledgements section.

## Acknowledgements

We wish to thank beta testers and users for feedback and patience throughout the ongoing development process of SuperFreq. This includes Aliaksei Holik, Göknur Giner, Jan Schröder, Christophe Lefevre, Gil Hornung, Kirill Tsyganov, Arun Ramani, and more.

The results here are based upon data generated by the TCGA Research Network: http://cancergenome.nih.gov/. The combination of donors, research teams and data sharing facilities at GDC provide an invaluable resource for cancer genomics research. We showed an example from a public diffuse large B-cell lymphoma dataset [phs000450.v2.p1], part of the Slim Initiative for Genomic Medicine (SIGMA), a joint U.S.-Mexico project funded by the Carlos Slim Health Institute. Another example was from the Epigenetic studies in Acute Myeloid Leukemia [phs001027], which was supported by K08CA169055 (F.E. Garrett-Bakelman), Starr Cancer Consortium I4-A442 (A.M. Melnick, R. Levine and C.E. Mason) and LLS SCOR 7006-13 (A.M. Melnick).

This work was supported by grants from the Australian National Health and Medical Research Council (www.nhmrc.gov.au) (Project Grant to IJM 1145912; Independent Research Institutes Infrastructure Support Scheme grant 9000220), the Cancer Council Victoria (www.cancervic.org.au) (grant-in-aid to IJM 1124178), a Victorian State Government Operational Infrastructure Support (OIS) grant; a Victorian Cancer Agency fellowship (to IJM) and a Felton Bequest to IJM. We also wish to acknowledge the generous support of Mr. Malcolm Broomhead who provided philanthropic support for the research. The funders had no role in study design, data collection and analysis, decision to publish, or preparation of the manuscript.

