## Supplementary Figures for "SuperFreq: Integrated mutation detection and clonal tracking in cancer"

### SuperFreq: Supplementary Figures

[1] Corresponding authors.

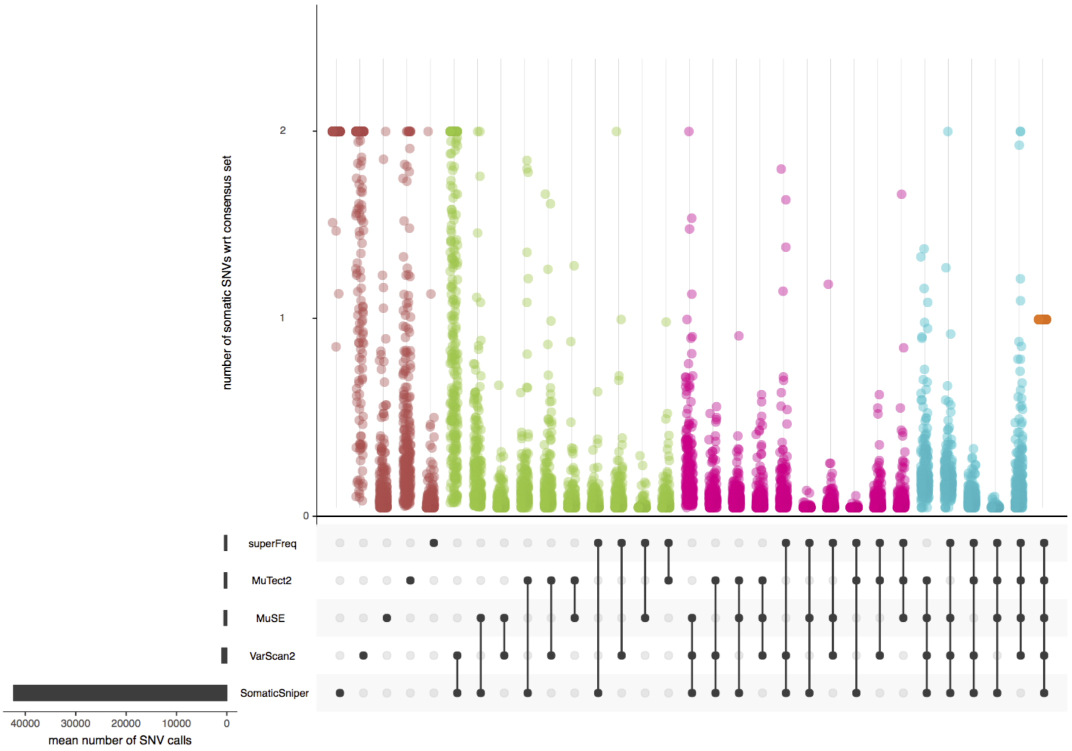


**Supplementary Figure 1**. Fraction SNVs called by subsets of 5 different SNV callers, relative to the number of SNVs called by all callers. Participants where less than 10 SNVs were called by all methods are not included. The fraction is capped at 2. Graphics produced with the help of UpSetR.

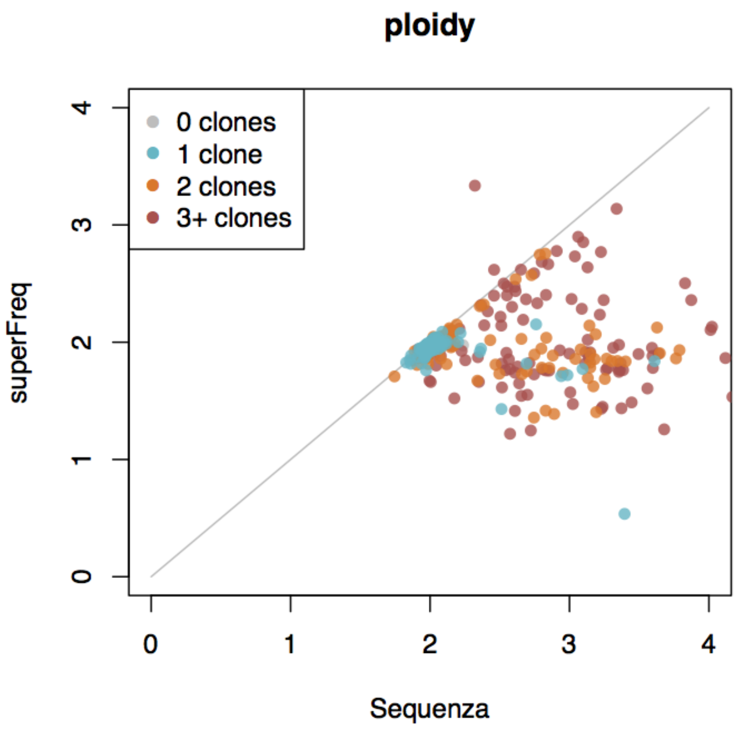


**Supplementary Figure 2**. Ploidy calls from Sequenza and superFreq, coloured by number of cancer clones called by superFreq.

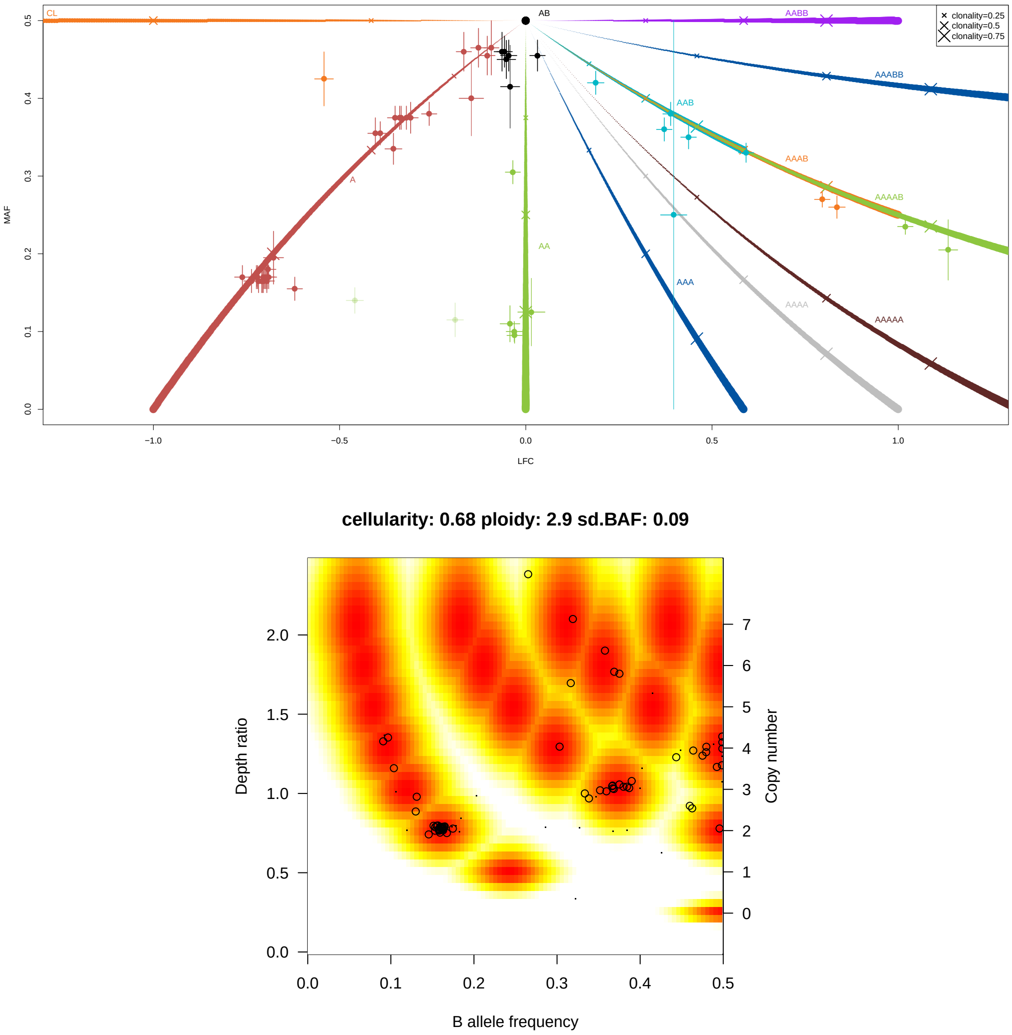


**Supplementary Figure 3**. Copy number calls and CNA model fits of TCGA-34-5240 (Lung Squamous Cell Carcinoma) from superFreq and and Sequenza. **a)** Maypole plot showing the model fit for the ploidy call (or equivalently LFC normalisation) in superFreq. Coloured lines show expected LFC and MAF of different copy number calls, with lines growing thicker with the clonality of the call. Dots show data from each segment with uncertainty in LFC and MAF, allowing for heteroscedasticity. Normalisation corresponds to a constant shift along the x-axis to make the crosses fit with the lines within errors. **b)** Sequenza model fit of the ploidy and purity call. Linear copy number call on the y-axis roughly corresponds to the x-axis of the maypole plot, and the x-axis in the Sequenza plot is the y-axis of the maypole plot. The single purity gives rise to single points for each copy number call, and the uncertainty is shown for the expected call rather than for the data points.

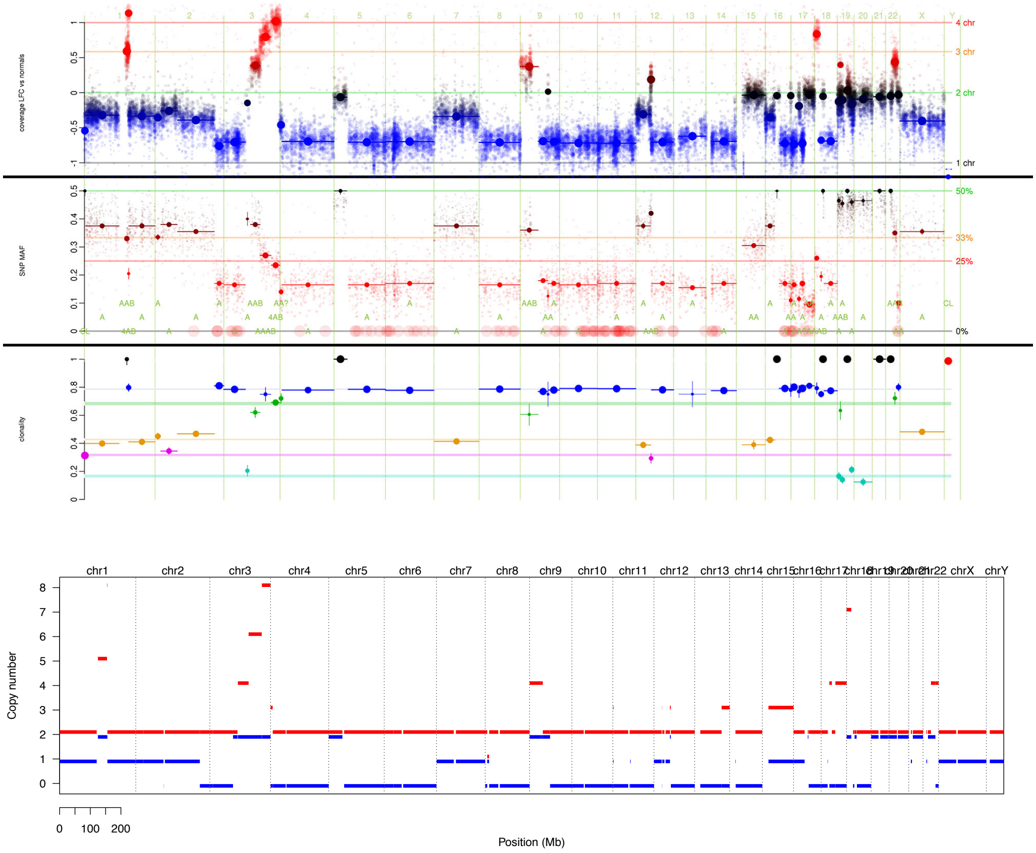


**Supplementary Figure 4.** Copy number calls and CNA model fits of TCGA-34-5240 (Lung Squamous Cell Carcinoma) from superFreq and and Sequenza. **a)** Copy number calls from superFreq, showing LFC, MAF and clonality of the call. The size of the dots represent accuracy, based on the adjusted limma estimates for LFC, and based on the effective coverage for the BAFs. Segments, shown as dots with horizontal lines, also shows error estimates through an error bar and point size, and the extension of the segment on the x-axis. CNA calls are shown below the BAF segments, where uncertain calls (inconsistent data) are marked with "?" or "??". **b)** Copy number profile from Sequenza with a purity of 0.68. Red shows major copy number, blue shows minor allele copy number.

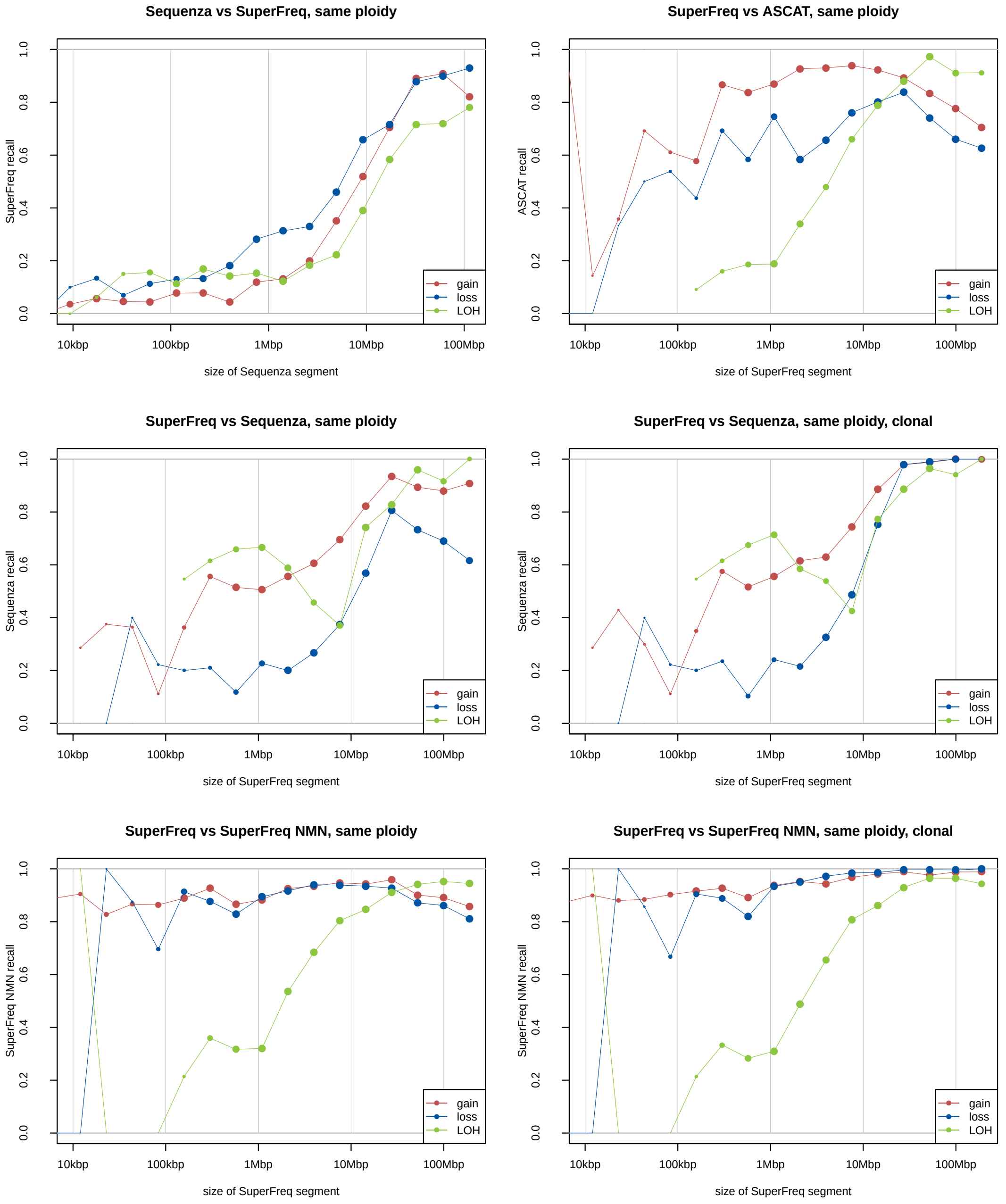


**Supplementary Figure 5**. Recall of gain, loss and CNN-LOH binned by size of the segment, limited to participants where the ploidy agrees within 0.2 between the methods. "Clonal" indicates that the truth segments are limited to CNAs where SuperFreq called a clonality above 0.5.

**a) CNA profiles**

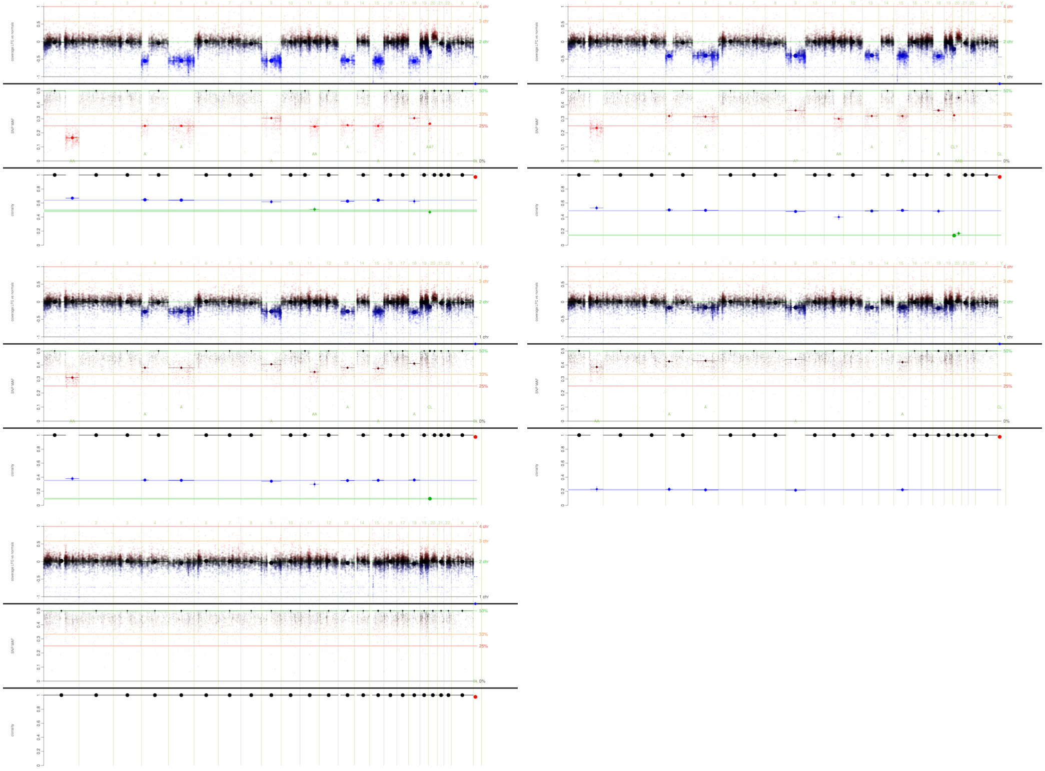

**b) River plot**

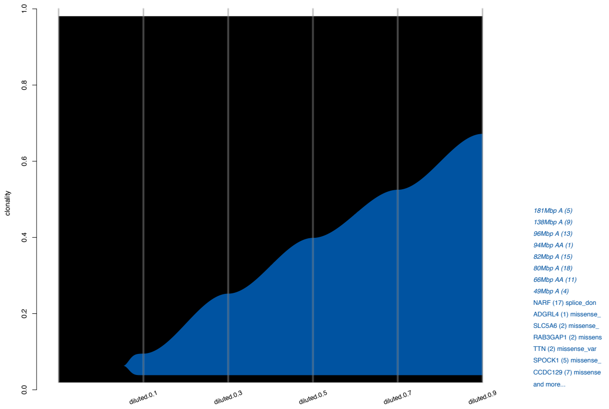


**c) Clonal tracking of CNAs**

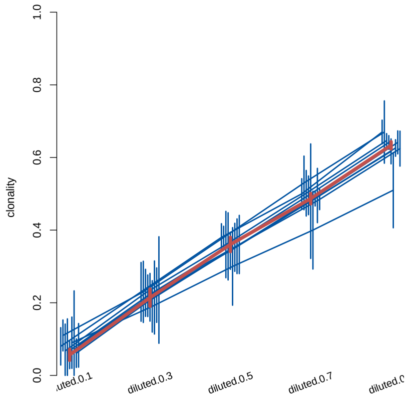


**Supplementary Figure 6**. Copy number calls and clonal tracking across the dilution series of TCGA-BQ-5879. a) Copy number calls at dilutions of 0.9, 0.7, 0.5, 0.3 and 0.1. b) River plot (germline variants removed) of the five dilutions analysed together. Although some CNAs are not called at the 0.3 dilution, and none is called at 0.1, they are still tracked and are assigned accurate clonalities as shown in c) where tracked CNAs are shown as blue lines, and the cancer clone is shown in red. SuperFreq shares the call and the segment coordinates across samples and queries the clonality by forcing the copy number call onto the segment in the other samples. In case the CNA is truly not present, the confidence interval is expected to overlap a clonality of 0. This analysis is performed without the matched normal sample, as the matched normal was used to dilute the cancer sample.

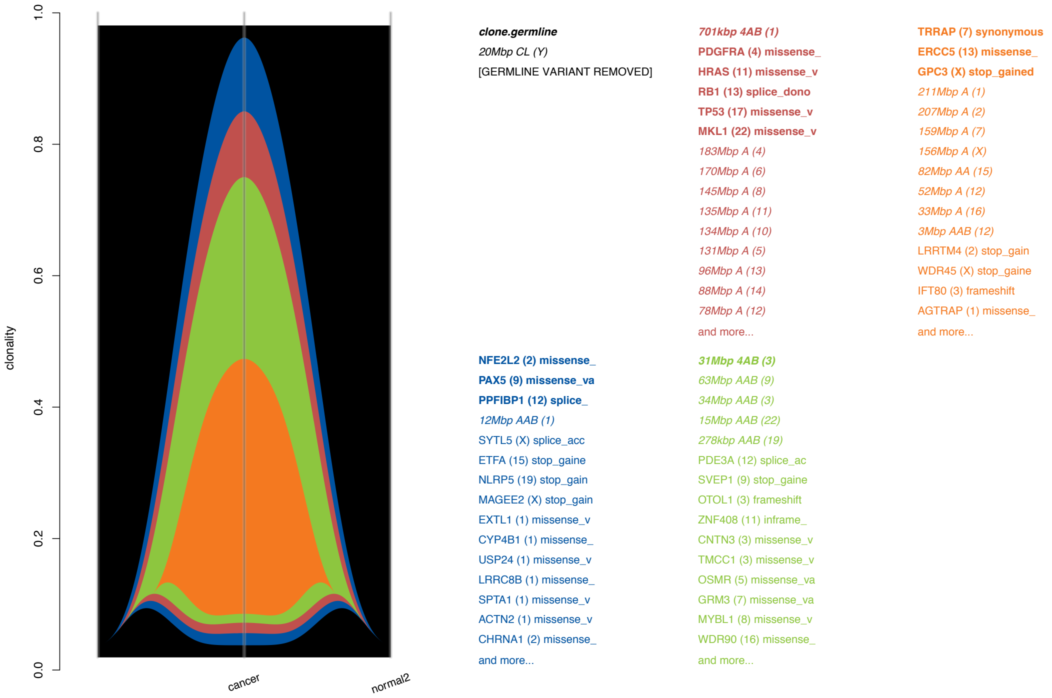


**Supplementary Figure 7**. Simulated clonal tracking of TCGA-34-5240 (Lung Squamous Cell Carcinoma). The original cancer has 4 clones called by superFreq. The copy number profile is shown in **Supplementary Figure 4**.

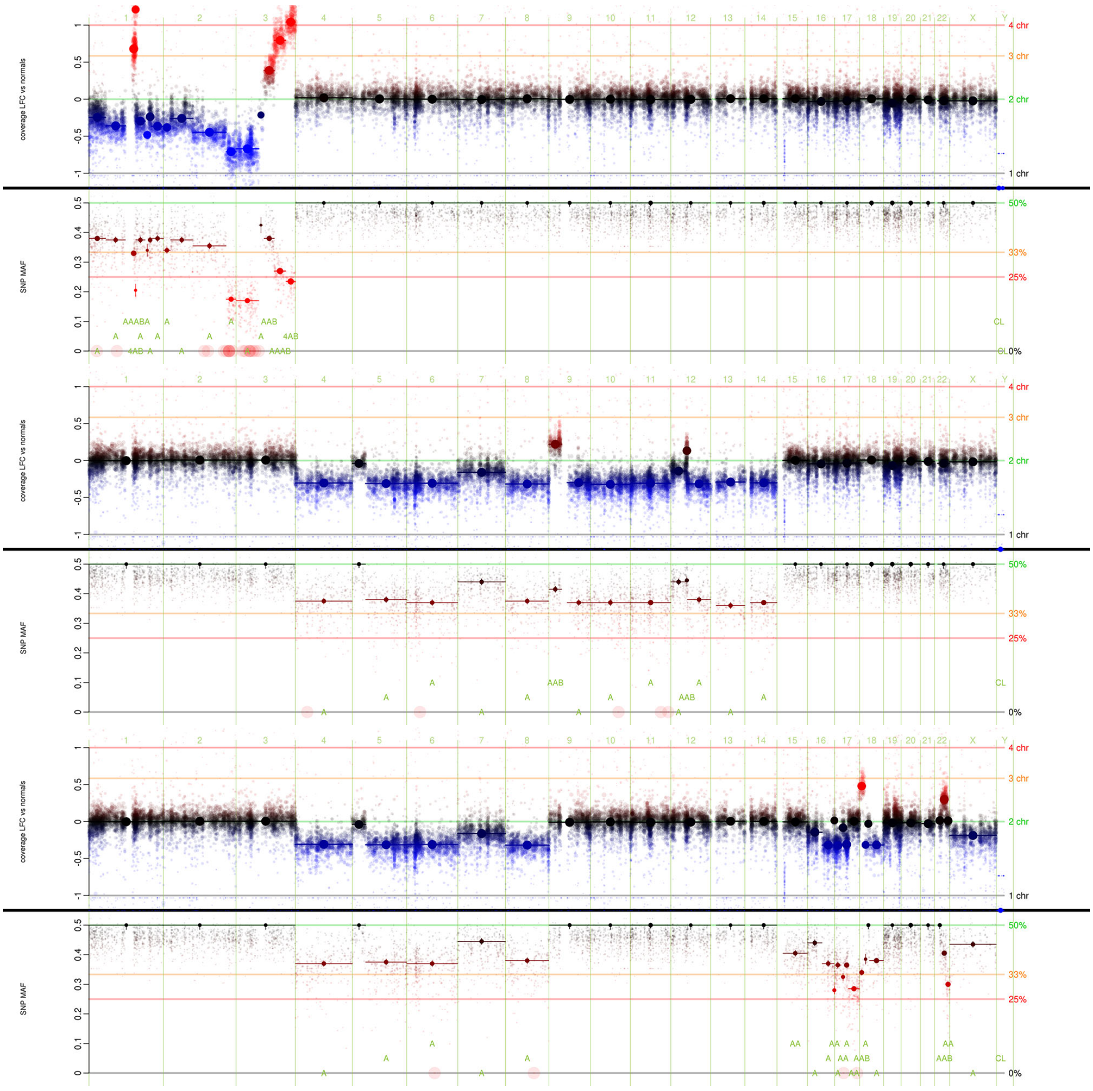


**Supplementary Figure 8**. Simulated clonal tracking of TCGA-34-5240 (Lung Squamous Cell Carcinoma). CNA calls over the genome showing LFC and BAF. The size of the dots represent accuracy, based on the adjusted limma estimates for LFC, and based on the effective coverage for the BAFs. Segments, shown as dots with horizontal lines, also shows error estimates through an error bar and point size, and the extension of the segment on the x-axis. CNA calls are shown below the BAF segments, where uncertain calls (inconsistent data) are marked with "?" or "??". The three simulated samples draw from mutations in different subsets of the chromosomes and of different admixtures of normal and cancer samples as illustrated by the copy number calls of the three samples. This process is described in **Figure 4a** in the main paper. The copy number profile of the original cancer is shown in **Supplementary Figure** 4.

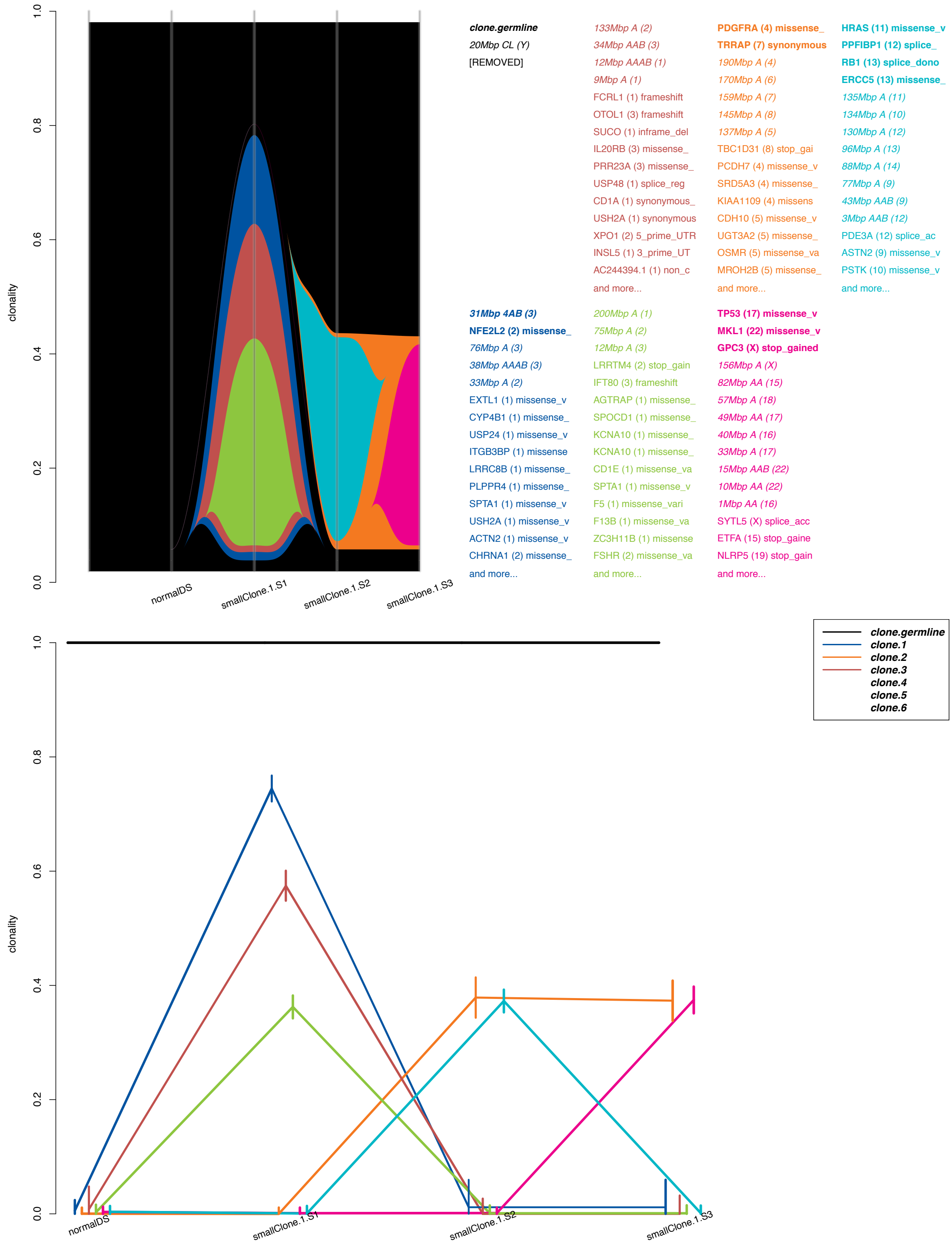


**Supplementary Figure 9**. Simulated clonal tracking of TCGA-34-5240 (Lung Squamous Cell Carcinoma). The superFreq clonal tracking of the simulated samples detects 3 subclones of the first clone based on the mutations on chr1 to chr3, while subclones are not detected for the other clones. We see that the mutations listed in each clone are found on the expected chromosomes from the schematic in **Figure 4a** in the main paper.

| | truth | TP | $FN_P$ | FP | $TP_g$ | $FP_g$ | $FP_{gP}$ |
|---|---|---|---|---|---|---|---|
| AML.080 | 296 | 273 | 23/23 | 230 | 271 | 58 | 30 |
| AML.084 | 10 | 8 | 2/2 | 263 | 5 | 88 | 55 |
| AML.102 | 9 | 7 | 2/2 | 216 | 3 | 76 | 47 |
| AML.110 | 13 | 12 | 1/1 | 193 | 5 | 48 | 26 |

**Supplementary Table 1**. truth: Total number of coding somatic variants called in SuperFreq with matched normal.
TP: Number of coding variants recalled (**T**rue **P**ositives) without a matched normal (total 91%).
$FN_P$: Fraction of lost coding variants (**F**alse **N**egatives) without matched normal that are present in **P**opulation databases dbSNP or ExAC (100%).
FP: Number of coding variants called without matched normal, not called with a matched normal (**F**alse **P**ositives).
$TP_g$: Number of variants recalled (**T**rue **P**ositives) without a matched normal after filtering on the ***g**ermlineLike* flag (total 87%).
$FP_g$: Number of coding variants called without a matched normal after germlineLike filter, not called with a matched normal.
$FP_{gP}$: Number of coding variants called without a matched normal after ***g**ermlineLike* filter, not called with a matched normal (**F**alse **P**ositives), that are present in **P**opulation databases dbSNP or ExAC.
We note that the number of false calls does not seem to depend on the number of true mutations, which confirms that the absolute number of false calls is a more robust measure of performance than normalised measures such as precision.

| AML.084 | Recall | FN in dbSNP | FP | runtime |
|---|---|---|---|---|
| 10 normals | 69/69 | 0/0 | 0 | 150m |
| 5 normals | 65/69 | 3/4 | 12 | 129m |
| 3 normals | 63/69 | 5/6 | 12 | 119m |
| 2 normals | 66/69 | 2/3 | 21 | 106m |

**Supplementary Table 2**. Recall, fraction false negatives in dbSNP, number of false positive somatic variants and runtime with 4 cpus in AML.084 with 5, 3 and 2 reference normals, using the analysis with 10 reference normals as truth. The runtimes are for the first run with the reference normals, subsequent runs of other samples using the same reference normals reuse gene counts and variants which decreases runtime. In our experience, the quality of the reference normals in mimicking the studied samples biases is more important than the number of reference normal samples.
