## Supplementary Methods for "SuperFreq: Integrated mutation detection and clonal tracking in cancer"

### SuperFreq: Supplementary Methods

[1] Corresponding authors.

#### Contents


**1 The superFreq algorithm** **1**
- 1.1 Error propagation 1
- 1.2 SNV filtering 2
- 1.3 Somatic SNV calling 3
- 1.4 Coverage analysis 4
  - 1.4.1 Variance consistency 5
- 1.5 Germline heterozygous SNPs 7
  - 1.5.1 Identifying germline heterozygous SNPs 7
  - 1.5.2 Statistical approach 8
  - 1.5.3 Reference bias correction 9
  - 1.5.4 Effective coverage 9
- 1.6 CNA calling 11
  - 1.6.1 Segmentation 11
  - 1.6.2 CNA calling and ploidy normalisation 11
  - 1.6.3 Allele Aware CNA tracking 13
- 1.7 Clonal tracking 17
  - 1.7.1 Clonality of somatic SNVs 17
  - 1.7.2 Clustering of mutations 17
  - 1.7.3 Non-repeating mutation assumption and convergent evolution 19

**2 TCGA analysis** **20**
- 2.1 Sample selection 20
- 2.2 Somatic SNV calls 20
- 2.3 Dilution and Slicing of the samples 20
- 2.4 Comparison of Copy Number Calls 21
- 2.5 SuperFreq Copy Number Sensitivity Assay 21
- 2.6 Clonal simulations 21


### 1 The superFreq algorithm

#### 1.1 Error propagation

Uncertainties in measurements are estimated and tracked throughout calculations in superFreq. Unless otherwise specified this is tracked as a single value interpretable as a confidence interval of 70%, roughly corresponding to a standard deviation of a normal distribution. While the first error estimate of observed values from the data (such as VAF or read depths) are done with different techniques adapted to the measurement, the propagation of the errors in subsequent calculation is generally consistent.

Unless otherwise specified, the uncertainty is propagated as a first order perturbation and combined as independent errors:

$$\Delta b = \sqrt{\sum_{i\in[1,N]} \left(\frac{\partial f}{\partial a_i}\Delta a_i\right)^2} \tag{1.1}$$

where $b = f(a_1, a_2, ..., a_N)$ is an observable with uncertainty $\Delta b$ calculated as function of $N$ previous observables $a_i$ with uncertainties $\Delta a_i$. When combining several observables into a consensus value, such as combining coverage LFCs in a copy number segment, or clonalities of individual mutations into the clonality of a clone, we use weighted mean with the inverse square error as weight. That is, if we want to find the consensus value $A$ from a set of independent measurement $a_i$ with errors $\Delta a_i$ we use

$$A = \frac{\sum_{i\in[1,N]} a_i/\Delta a_i^2}{\sum_{i\in[1,N]} 1/\Delta a_i^2} \tag{1.2}$$

and from eq. (1.1) it follows that

$$\Delta A = \frac{1}{\sqrt{\sum_i 1/\Delta a_i^2}}. \tag{1.3}$$

This allows us to carry the uncertainty of each measured VAF and read count all the way through CNA calling, somatic SNV calling and clonal tracking. The framework allows us to incorporate biological, theoretical or other systematic error sources where needed. We don't make any explicit assumptions on the distribution of the uncertainty, in particular we do not claim that they are Gaussian. This does not completely bypass the problem of choosing parameterisation of the uncertainty distribution though. For practical purposes, when calling whether an observable is significantly different from another, we set a limit in number of errors difference, typically 3. This replaces the choice of parameterisation and cut on probability or p-value. For comparison, cutting at three error bars corresponds to a p-value cut of 0.3% for a normal distribution, or a 3% cut for a t-distribution with 5 degrees of freedom.

##### 1.2 SNV filtering

Only the genomic positions from the supplied VCF files are utilised. Any variants more than 300bp away from a capture region are discarded. Positions are summarised for each individual, and the pileups for all positions indicated in the VCF files are imported from the BAM of all samples of that individual, as well as the reference normals. SuperFreq checks for a range of quality issues and assigns the following flags:

- "Bq", **Base Quality** is triggered if the variant reads have a significantly lower base quality than the reference reads ($p_{bq} < 0.01$, Mann-Whitney U-test) and the mean base quality is at least 10 lower. This flag is also used if the overall mean base quality is below 20, or strictly less than 10% of the variant reads achieve a base quality of 30.

- "Mq", **Mapping Quality** is triggered if the variant reads have a significantly lower mapping quality than the reference reads ($p_{mq} < 0.01$, Mann-Whitney U-test) and the mean mapping quality is at least 10 lower. This flag is also used if the overall mean mapping quality is below 20, or strictly less than 10% of the variant reads achieve a mapping quality of 30.

- "Sb", **Strand Bias** is triggered if the variant reads have a significantly different strand ratio than the reference reads ($p < 0.001$, Fisher's exact test). In our experience, strand bias is rarely the only warning sign of a false call. The lower prior expectation for this flag manifests in a reduced cut for the p-value.

The variants are also compared to the reference normal samples, and a set of flags are assigned to variants that are present in the normals:

- "Nnc" and "Nnn", **Normal Noise Consistent or Non-consistent** triggers if the variant is present at more than 10% in any of the normal samples. Consistency is determined based on whether all normal samples are consistent with the same background frequency (fisher's exact test, $p > 0.01$). Variants in dbSNP are allowed to have frequencies consistent with 0.5 or 1 as well without being flagged.

- "Mc", **Many Copies**. Variants where the reference normals have more than 10 times the median coverage, with the median across variants. This is often associated with regions that are present multiple times in the human genome, but only present once in the reference, leading to inflated coverage and heterozygous germline variants deviating from 50%. This flag is disabled when running in RNA mode, as highly varying coverage is expected from varying expression between genes.

By using the reference normals in this manner, superFreq does not need to blacklist or mask repeat or low complexity regions.

##### 1.3 Somatic SNV calling

SuperFreq calls somatic variants by comparing to the match normal if present, or based on population frequencies and clonal tracking otherwise.

To be called as a somatic variant when a matched normal is present, a variant has to pass a number of filters, and the somatic score $s$ starting at 1 can receive multiplicative penalties for quality issues.

- Number of variant reads in the normal $V_n$ has to be less than $\sqrt{C_n}/2$, where $C_n$ is the coverage in the normal. Non-zero normal variant counts get a penalty factor to the somatic score of $(1 - 5V_n/C_n)$. This requires a close to zero frequency in the normal, while still allowing the occasional miscalled base, especially at lower coverage.

- The sample has to have a significantly larger frequency than the normal (FET, p $<$ 0.01) and the sample frequency has to be at least 0.05 larger than the normal. The variant is assigned a penalty of $1 - 100p_{\text{FET}}$ where $p_{\text{FET}}$ is the fisher exact test p value.

For samples without matched normal, the requirements are:

- Each variant is looked up in dbSNP and ExAC and classified as one of
  - Not present.
  - Present, but at unkown population frequency.
  - Rare: present at population frequency below 0.001.
  - Common: present at population frequency equal to or above 0.001.

  Variants have to be either rare or not present in both databases, or rare in one database and unknown in the other. Here unknown frequencies paired with not present in the other database are not called as somatic, as it can be a common variant that is not called in the other data base due to read coverage or limited sensitivity for other reasons.

Some requirements are shared independently of whether a matched normal is found or not.

- Each of the Bq, Mq and Sb p-values in section 1.2 are multiple hypothesis corrected to false discovery rates with the Benjamini-Hochberg procedure and multiplied to the somatic score, with a lower limit of 0.8. This is not intended as filtering, as only unflagged variants are considered, but will decrease the confidence in the call if the quality scores are suspicious.
- If the coverage over the variant in the sample is below 10 reads, a penalty of $C_s/10$ is multiplied to the score, where $C_s$ is the coverage in the sample.
- The variant has to have a significantly larger frequency than the reference normals. The somatic score gets a penalty of $1 - p_r$ where $p_r$ is the p value from a binomial test of the samples variant read count and coverage against the total frequency over the reference normals. A penalty of $1 - 5f_r$, where $f_r$ is the total frequency in the reference normals, is also multiplied to the score. These penalties are mainly aimed at the case of no matched normals, where they catch low frequency noise that passed filters, or common germline variants with low frequencies due to multiple copies of the region over the genome.

If the sample is marked as normal, then somatics are called as if no matched normal is present, even if a second normal sample is available for the individual, in effect identifying rare germline variants. These will be tracked in the clonal analysis and can show up in the germline clone in the output.

In the case of multiple matched normal samples, the read counts are merged when compared to the non-normal samples.

##### 1.4 Coverage analysis

The coverage of each sample is compared to the coverage of the reference normal samples, similarly to a standard differential expression analysis with limma-voom. First, fragments are counted with FeatureCounts over each of the padded (300bp on each side) capture regions for all samples, including the reference normals. The counts are then corrected for:

- **GC content**. A loess curve is fitted to $\log(N_{rs}/L_r)$ as function of GC content over the region $r$ for each sample $s$, where $N_{rs}$ is the number of reads over the region $r$ in sample $i$, and $L_r$ is the length of the capture region $r$. The loess fit is weighted by $\sqrt{\langle N_{rs}\rangle_s}$ with $s$ running over all samples. The counts are then divided by the value of the weighted loess fit, maintaining total read count. This is illustrated in **Supplementary Methods Figure 1**.

- **MA-bias** by capture region. The log fold change $M_r = \ln_2(N_{rs}/N_r)$, where $N_r$ is the mean count over the reference normal samples, are plotted against $A_r = \ln_2(N_{rs}N_r)/2$. A loess fit is made to the curve, and the counts of each region $r$ are corrected with the inverse of the loess fit for that $A_r$. This is illustrated in **Supplementary Methods Figure 2**.

- The reference normal samples are **sex corrected** to two copies of every chromosome, meaning that male samples have their X and Y chromosome counts doubled while female sample maintain their X chromosome counts but have their Y chromosome counts removed from the analysis. The sex of a reference normal sample $s$ is set to female if the coverage on the X chromosome is at least 10 times the coverage of the Y chromosome: $\sum_{r\in X}(N_{rs})/\sum_{r\in X}(L_r) > 10\sum_{r\in Y}(N_{rs})/\sum_{r\in Y}(L_r)$, otherwise it is set male. $X$ and $Y$ are the sets of capture regions in the X and Y chromosome respectively.

The corrected counts are summarised by gene and analysed for differential coverage using limma-voom with sample weights. Analysis is done with each sample compared to the reference normals with a one-against-many design matrix. The output is a log fold change $\beta_{gs}$ for each sample $s$ and gene $g$ with respect to the reference normals. The moderated t-statistic $t_{gs}$ from Limma is used for an error estimate $\Delta\beta_{gs} = |\beta_{gs}|/t_{gs}$ together with the posterior degrees of freedom $d$. In downstream analysis $\beta_{gs}$ and $\Delta\beta_{gs}$ are propagated according to eqs. (1.1, 1.2, 1.3), but when we calculate the likelihood $p_{12}$ that two regions 1 and 2 have the same $\beta$ during copy number segmentation, we utilise the t-distribution from limma with

$$p_{12} = t\left(\frac{|\beta_1 - \beta_2|}{\sqrt{\Delta\beta_1^2 + \Delta\beta_2^2}}, d\right), \tag{1.4}$$

where $t(\cdot,\cdot)$ is the t-distribution. In this way we can use both the degree of freedom estimated by limma-voom, as well as using the dynamic error propagation when combining measurements.

##### 1.4.1 Variance consistency

Limma-voom provides excellent variance estimates for RNA-seq, taking full advantage of the available information. For the coverage analysis of DNA of exome-like data, we have an additional piece of information though: we expect the vast majority of neighbouring capture regions to have the same copy number. We can use this information to assess the limma-voom variance estimates on a by-sample basis, and correct when the variance

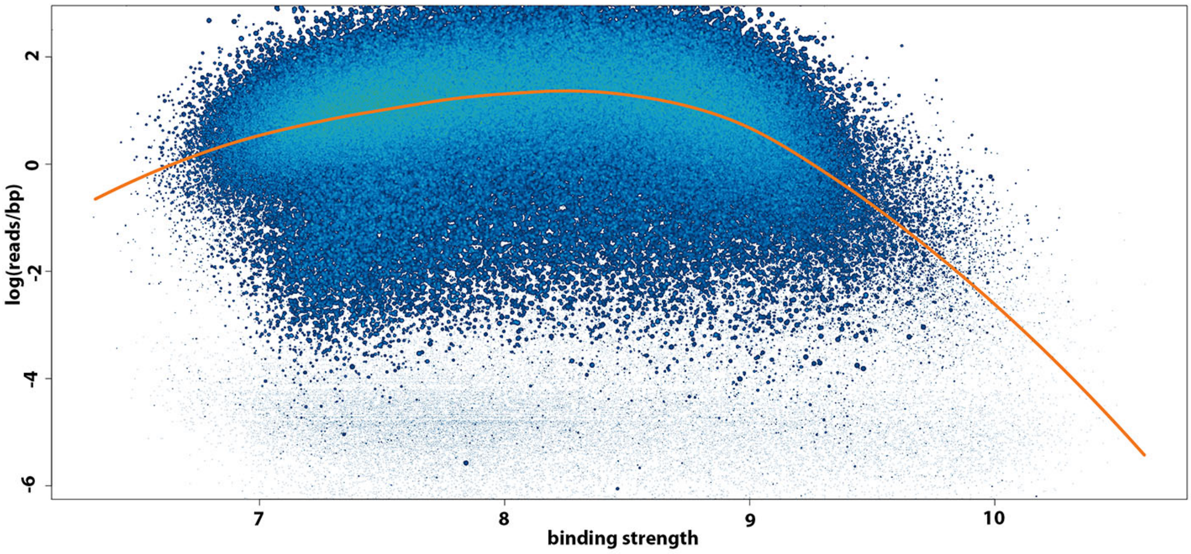


**Supplementary Methods Figure 1.** Log read depth as function of DNA binding strength (which is closely correlated to GC content), by capture Region. The orange line shows the loess fit that is used to correct the counts. Data from TCGA-A3-3320.

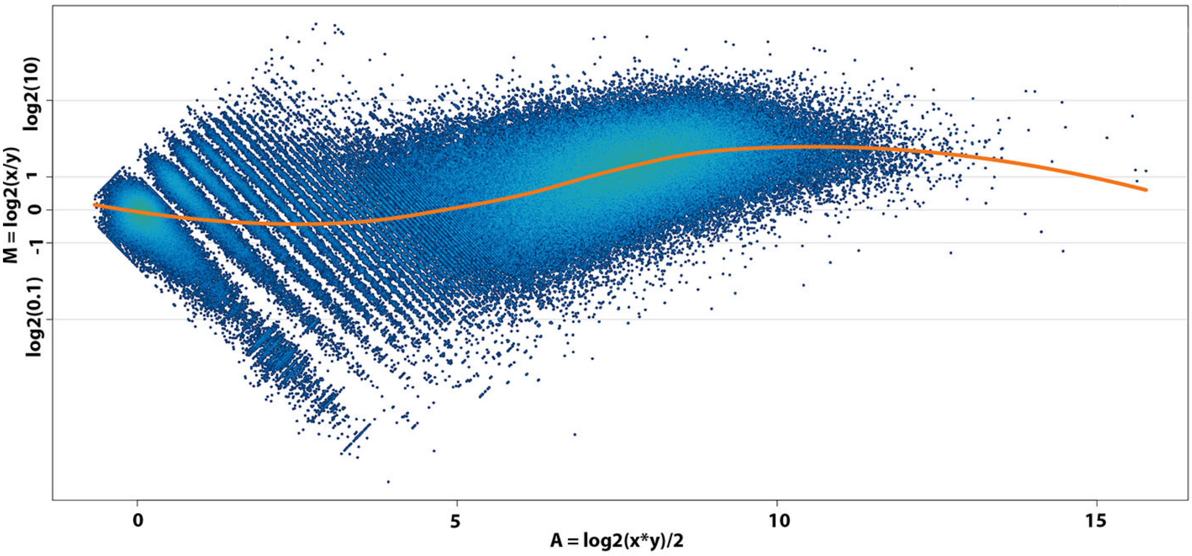


**Supplementary Methods Figure 2.** MA plot of the smeared read counts by capture region, compared to the reference normals. The orange line shows the loess fit that is used to correct the counts. Data from TCGA-A3-3320.

estimates do not describe what we see in the data. To do this, superFreq compares the difference in LFC between neighbouring capture regions divided by the error, to what is expected from the difference of two t-distributions with $d$ degrees of freedom. To remain robust to outliers, such as true copy number breakpoints, we restrict the measure to the median of the distribution. If the measured median of the difference is larger than the median of the expected distribution, superFreq adds a constant number to all variance estimates so that the measured median matches the expected value. SuperFreq does not decrease overestimated variance estimates. This correction, a one-parameter fit, often yields a surprisingly good fit between the measured and expected distribution of differences, as is illustrated in **Supplementary Methods Figure 3**.

#### 1.5 Germline heterozygous SNPs

##### 1.5.1 Identifying germline heterozygous SNPs

Germline heterozygous SNPs are identified from the SNVs that passed the quality filters. We require a population frequency larger than 1% in dbSNP and ExAC, and use the VAF to identify heterozygous SNPs. If a matched normal is present, we require a VAF between 35% and 65% in the normal sample, and a p-value (binomial against 50% VAF) above 10%. If a matched normal is not present, we have to accomodate heterozygous SNPs that are affected by CNAs and deviate from 50% VAF. Close to 100% we risk picking up homozygous SNPs, and at low frequencies there is an increasing amount of noise. As a compromise we take SNPs between 5% and 95%. In theory this does not allow us to detect LOH with clonalities close to 100%, but in practice some noise variants are usually picked up just above 5%

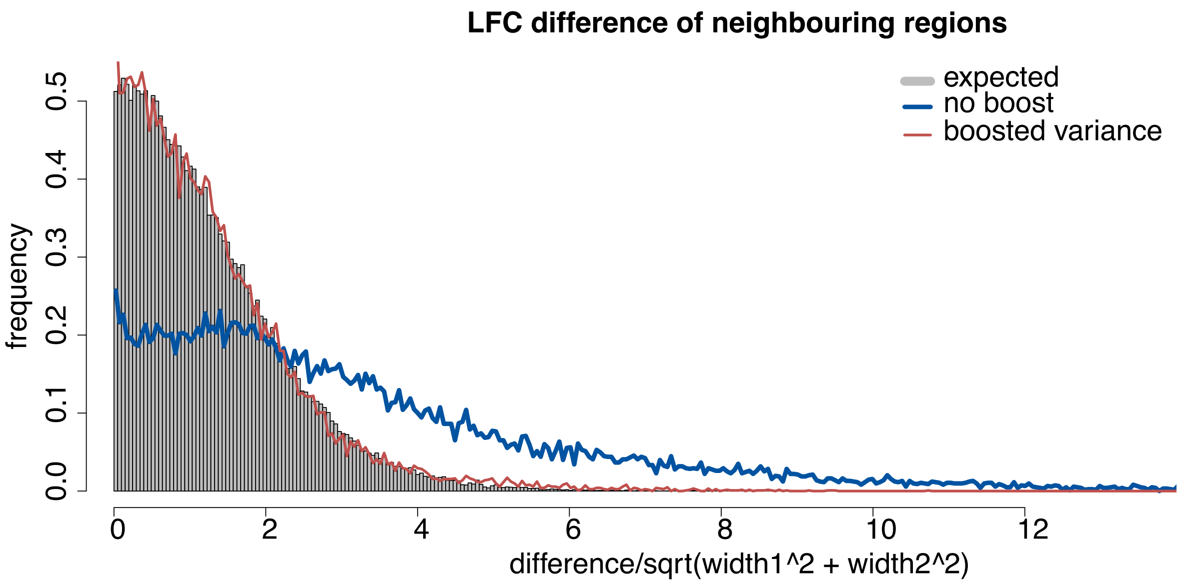


**Supplementary Methods Figure 3**. The difference in LFC between neighbouring capture regions, divided by the joint uncertainty. Grey bars show the theoretical expected distribution, and the blue line shows the distribution from the raw limma-voom variance estimates. Red line shows the distribution when all LFC variances are increased by a constant to match median of the distributions. In this and many other cases, we see a surprisingly good match of the grey and red distribution considering that it is a one-parameter fit. Data from TCGA-A3-3320.

which will nonetheless give a high clonality LOH call in the absence of true heterozygous SNPs.

The search for heterozygous germline SNP is run unchanged on the sex chromosomes, but are ignored on the Y chromosome and on the X chromosome of males. The number of heterozygous germline variant calls in these regions are counted and used for a naive extrapolation to the total number of false calls genomewide.

##### 1.5.2 Statistical approach

It is not trivial to detect small deviations from 0.5 in the germline heterozygous SNP frequency. As the SNP frequencies are expected to go both up and down from 0.5, it is not sufficient to look for a deviation in mean frequency. To solve this, all frequencies are mirrored around 0.5 down to the smaller frequency of the reference and variant frequencies, referred to as the minor allele frequency $f$. A copy number deviation now shows up as a shift in mean, but on the other hand the absence of copy number alterations is no longer associated with a mean of 0.5, making the null hypothesis more complex. The heart of the problem is that high variance in the SNP frequency is giving a very similar signal as a subclonal copy number change, both slightly broadening the frequency distribution around 0.5. The problem is compounded by the presence of incorrectly called germline heterozygous SNPs, especially in the absence of a matched normal, that can further increase variance.

The algorithm in superFreq that determines if a CNA region is deviating from the null hypothesis of 50% allelic balance can be summarised in four steps:

- **Find the best guess for an alternative frequency.** All frequencies $f$ between 0 and 0.5 are scanned in steps of 0.1%, and the likelihood $p_i$ for each SNP $i$ to come from the frequency $f$ or $1-f$ is calculated with binomial distributions. The product over all the SNPs in the region $\prod_i p_i$ represent the likelihood of the SNPs having a frequency $f$. The frequency $f_a$ with the maximum likelihood is selected as the alternative hypothesis. Note that due to natural variation of the frequencies (due to finite coverage and other sources), this alternative hypothesis is lower than 0.5 even for truly null regions, typically between 0.4 and 0.45 for standard exomes.

- **Filter SNPs that are not consistent with null or alternative frequency** Any SNP $i$ that has a $p_i$ below 0.05 in both the null ($p_{i,\mathrm{null}}$) and alternative ($p_{i,\mathrm{a}}$) hypothesis are removed. This is meant to filter out false SNP calls that mostly will not have frequencies consistent with null or alternative frequencies. This means that 5% of the true SNPs will be removed as well, meaning a small loss in power, but we have found the trade off to be beneficial.

- **Calculate mean log likelihood ratio (MLLR)** Each SNP $i$ now is assigned a null likelihood distribution (binomial around 0.5) and an alternative likelihood distribution (superposition of two binomials at $f_a$ and $1-f_a$), all binomial with the read depth of the SNP $i$. A log likelihood ratio (LLR) $l_i = \log(p_{i,\mathrm{null}}/p_{i,\mathrm{a}})$ is calculated from the two distributions, and the MLLR is taken over all SNPs: $L = \langle l_i \rangle_i$.

- **Compare to expected MLLR from null or alternative hypothesis** We now first assume that the null hypothesis is true, and calculate the expected value $L_{\text{null}}$ of the statistic $L$, as well as its variance $V_{\text{null}}$. We then do the same, assuming that the alternative hypothesis is true, giving the mean $L_a$ and variance $V_a$ of the statistic. These statistics are shown in the diagnostic plots, in the directory called MAFstats, shown in **Supplementary Methods Figure 4**. We assume normal distributions of $L$, which is accurate for many SNPs from central limit theorem, with a width of $V$, and calculate the likelihood of the region being consistent with the null $p_{\text{null}}$ or alternative $p_{\text{a}}$ hypothesis. $p_{\text{null}}$ is capped at a minimum $10^{10L}$ for negative $L$, in order to limit too strong null rejection from a very small average LLR from many SNPs. The posterior probability of the region being 50% heterozygous is $p = p_{\text{null}}/(p_{\text{null}} + p_{\text{a}})$ where we have assigned equal prior probability to both null and alternative hypothesis.

###### 1.5.3 Reference bias correction

To compensate for reference bias, we estimate the bias from the heterozygous SNPs in the reference normal samples. We extract clean dbSNP variants that are close to 0.5 (between 0.2 and 0.8, and a binomial p-value above 1%) and calculate the weighted mean of the variant frequency $F = \sum_i v_i / \sum_i c_i$ where $i$ runs over variants, $v_i$ is the variant count and $c_i$ is the coverage. The variant loss, the ratio of variant reads that are lost is now $L = 1 - F/(1 - F)$. All frequencies are adjusted by this average loss to a new frequency

$$f' = \frac{f}{f + (1 - f)(1 - L)}.$$

The process is repeated with the corrected frequencies until equilibrium is reached when $L$ changes less than $10^{-5}$ between two iterations. This correction is applied to the heterozygous SNPs throughout the CNA analysis, but is not used for the somatic SNVs. If $L$ is unrealistically small or large, or the iteration does not converge, reference bias correction is not carried out, corresponding to $L = 0$.

###### 1.5.4 Effective coverage

SNP frequencies are affected by multiple sources of variance, and a binomial assumption on the frequency distribution is often not capturing this properly, especially not for high coverage SNPs. There are multiple alternative distributions with an extra parameter that maintains an approximately binomial behaviour at low coverage but maintain a minimum variance at high coverage. SuperFreq instead transforms the coverage $c$ to an effective coverage

$$c_e = \frac{c(1 + c/C_m)}{1 + c/C_m + c^2/C_m^2} \tag{1.5}$$

where $C_m$ sets the maximum effective coverage that $c_e$ converges to for large raw coverage $c$. $C_m$ is a superFreq parameter `maximumCoverage` set to 150 by default. This monotonous parameterization scales as the raw coverage $c$ to first order for low coverage $c \ll C_m$, and converges to a constant $C_m$ for $c \gg C_m$.

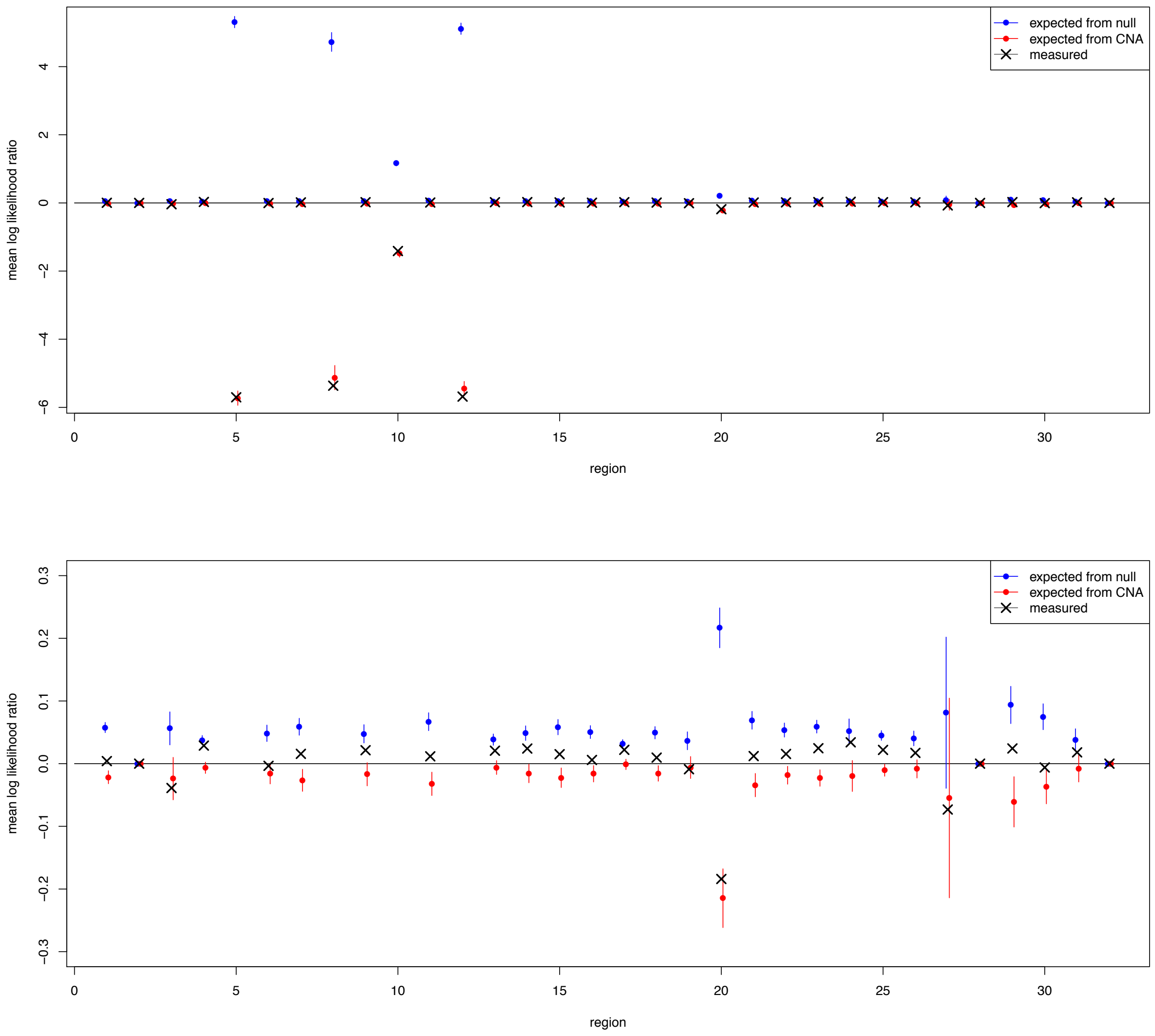


**Supplementary Methods Figure 4.** Statistics on the heterozygous SNPs. x-axis runs over segmented regions. Red dots show the expected outcome from the null hypothesis $L_{\text{null}}$ with $V_{\text{null}}$ as error bars, and similarly blue blue dots show the expected outcome from the alternative hypothesis $L_a$ with $V_a$ as error bars. Black dots show actual data. Second panel is zoomed in on the y-axis. Data from TCGA-A3-3320.

The variant count is adjust by the same factor to maintain the VAF, and all downstream statistics on the germline SNPs for the copy number analysis is done with the effective coverage and variant counts.

##### 1.6 CNA calling

###### 1.6.1 Segmentation

Segmentation is done by hierarchical clustering of neighbouring regions on each chromosome, starting with each gene as a separate region. The probability that two neighbouring regions share both minor allele frequency $f$ and coverage LFC $\beta$ consists of a contribution from the $f$, a contribution from $\beta$, and a prior probability based on one expected CNA breakpoint every 100Mbp.

A consensus frequency $f$ over both regions is calculated from the summed coverage and variant counts across all SNPs in both regions. Then the coverage and variant counts are summed for each region separately and each is tested against the consensus frequency $f$ with a binomial test. The two p-values are combined using Simes procedure into a likelihood $p_f$ that both regions share SNP frequency $f$. Fisher's exact test would perhaps be more appropriate for this test, but is computationally expensive for large numbers of SNPs and the current implementation seems to perform similarly enough in the relevant range.

We also calculate the likelihood of both regions coming from a 50% heterozygous state by combining the posterior probabilities from section 1.5.2 with Simes test into $p_{f_0}$. The largest of $p_f$ and $p_{f_0}$, referred to as just $p_f$ below, is used.

The two regions are compared for coverage, and we get a likelihood $p_\beta$ that both regions share coverage calculated from eq. 1.4.

$p_f$ and $p_\beta$ are combined to a likelihood $p_{f\beta}$ that the regions share both $f$ and $\beta$ using Fisher's method of combining p-values. Finally we use a prior $p_p = e^{-dx/1e8}$, where $dx$ is the distance between the two regions, to calculate the posterior probability

$$p = \frac{p_p p_{f\beta}}{p_p p_{f\beta} + (1 - p_p)}$$

of no copy number break between the regions, where we assume that the data is always consistent with a non-zero (but possibly small) difference in $\beta$ or $f$.

The neighbours with the largest $p$ are merged into one region, and the score is recalculated for the new neighbours. This cycle is repeated until no neighbouring segments have Probability $p$ larger than 0.05 to be paired.

###### 1.6.2 CNA calling and ploidy normalisation

After segmentation, the coverage data is normalised to correspond to ploidy, and absolute copy numbers are called. Each region has a $\beta$ and $f$ with corresponding uncertainties that are matched against possible absolute copy number events of any clonality, assuming a heterozygous diploid (AB genotype) background. This is displayed in the maypole plot in **Supplementary Methods Figure 5a**.

To normalise the coverage data, equivalent to determining the ploidy of the sample, a shift $s$ is added to $\beta$ of all regions. $s$ is selected so that as many regions as possible are consistent with a copy number call at some clonality within uncertainties. This is done by first sampling a wide range of $s$ to calculate a fitness score and then fine tune by sampling at smaller interval close to the best scores. The fitness score $F(s)$ for a shift $s$ is

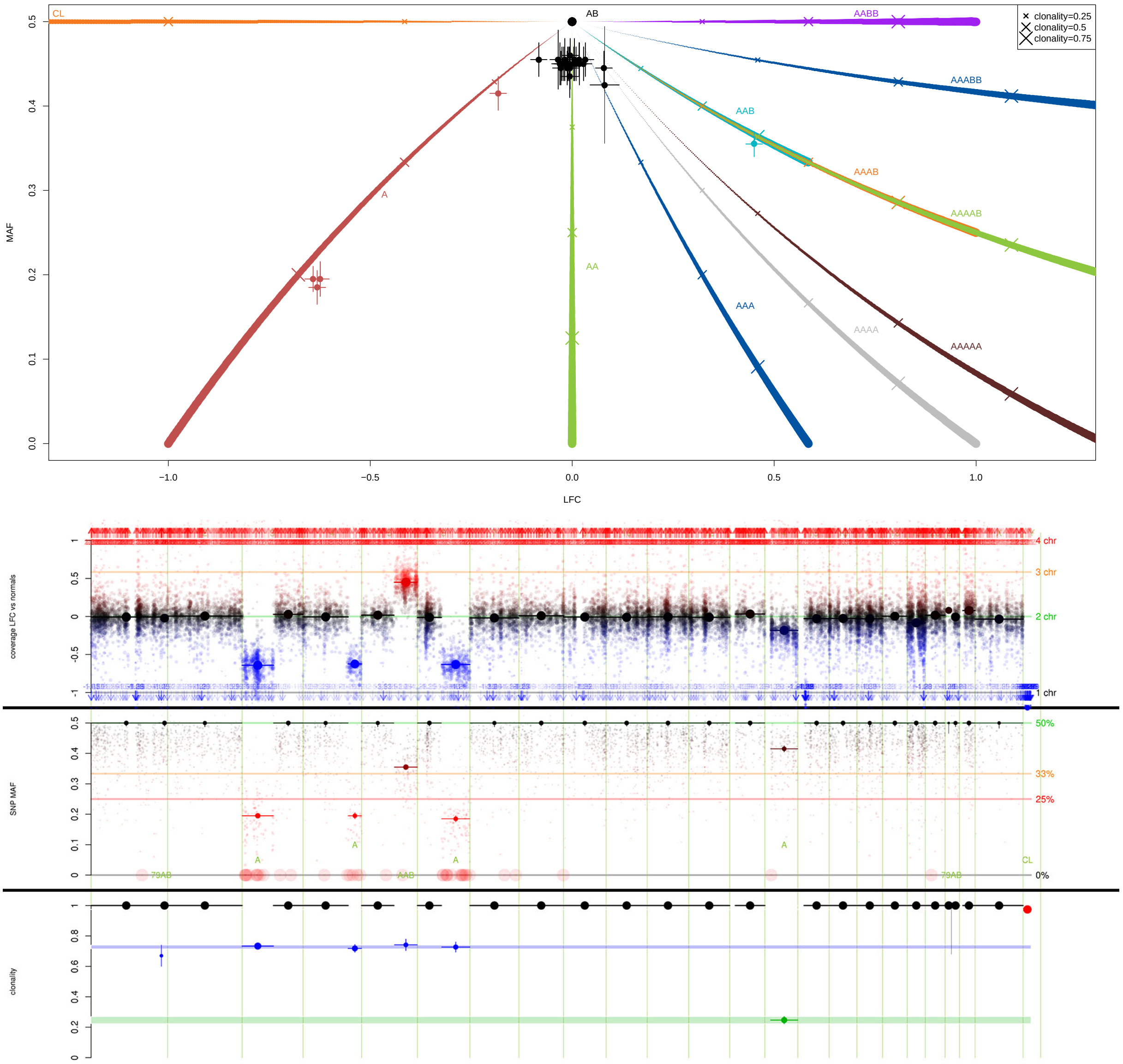


**Supplementary Methods Figure 5**. Top panel: Maypole plot showing the LFC with respect to the reference normals on the x-axis and the minor allele frequency on the y-axis. Each dot is a segment, and error bars show the uncertainty. Bottom panel: CNA calls over the genome of the same sample as in the top panel, showing LFC, MAF and clonality of the call. The size of the dots represent accuracy, based on the adjusted limma estimates for LFC, and based on the effective coverage for the BAFs. Segments, shown as dots with horizontal lines, also shows error estimates through an error bar and point size, and the extension of the segment on the x-axis. CNA calls are shown below the BAF segments, where uncertain calls (inconsistent data) are marked with "?" or "??". Data from TCGA-A3-3320.

$$F(s)^2 = \frac{1}{N}\sum_i \min_{n,c}\left(\sigma_{n,c,i}(s), 3\right)^2$$

| call | AB | A | AA | AAA | AAAA | AAAAA | AAB | AAAB | 4AB | 5AB |
|---|---|---|---|---|---|---|---|---|---|---|
| prior | 10 | 1 | 1/2 | 1/3 | 1/4 | 1/5 | 1 | 1/2 | 1/3 | 1/4 |

| call | 6AB | 9AB | 19AB | 39AB | 79AB | AABB | AAABB | 4ABB | CL |
|---|---|---|---|---|---|---|---|---|---|
| prior | 1/5 | 1/10 | 1/10 | 1/10 | 1/10 | 1/2 | 1/4 | 1/5 | 1/2 |

**Supplementary Methods Table 1**. The allowed CNA calls and their relative priors in superFreq. NAB means N copies of the A allele, and one copy of the B allele. CL means complete loss of both alleles. The priors are the inverse of the edit distance from AB, restricted to the range $[0.1, 10]$. The above scores are first normalised to $p_n$ so that they sum to 1, and $\sigma_n$ is the absolute value of the $p_n/2$ quantile of the normal distribution.

$$\sigma_{n,c,i}(s) = \sigma_n + \sqrt{\left(\frac{(\beta_i + s) - \beta_{n,c}}{2\Delta\beta_i}\right)^2 + \left(\frac{f_i - f_{n,c}}{2\Delta f_i}\right)^2}$$

where $i$ runs over all $N$ segments, $n$ runs over all copy number genotypes and $c$ runs over clonalities. $\sigma_n$ is a prior penalty for each CNA call $n$, based on the number of added or removed alleles as shown in table 1. $F(s)$ can be understood as the RMS of the number of uncertainties to the closest copy number call, with a maximum of 3 for segments that don't fit anything well.

With the normalisation shift $s_0$ fixed so that $F(s_0)$ is a global minimum, each segment $i$ is assigned a copy number call $n$ and clonality $c$ that minimises $\sigma_n + \sigma_{n,c,i}(s_0)$. The normalisation process can be thought of as shifting the x-axis in the top panel of **Supplementary Methods Figure 5**. To contrast, a high ploidy sample is shown in **Supplementary Methods Figure 6**.

**Postprocessing CNA calls** The copy number calls are then post processed for artifacts. To avoid segments that are based on false low frequency SNP calls, especially prevalent when called without a matched normal, we merge neighbouring segments that are separated due to only a few SNPs. We require that the regions have consistent $\beta$ within an error, and that one of the regions have either all of its heterozygous SNPs within a region of 100kbp, or an inconsistent $\beta_i$ and $f_i$ with any copy number call: $\sigma_{n,c,i}(s_0) > 2$. We also merge neighbouring segments that have the same copy number call $n$ and that have clonalities $c$ within two uncertainties. If any segments are merged, the normalisation process for $s_0$ is repeated, until the postprocessing merges no additional segments.

##### 1.6.3 Allele Aware CNA tracking

During the tracking of CNAs across samples, superFreq also compares the direction of deviation from 50% in B-allele frequency (BAF) of the germline heterozygous SNPs. In case different alleles are affected by the CNA in different (such as gain of the "A" or "B" allele, giving an AAB or ABB genotype respectively), then the BAFs will deviate in opposite directions. When superFreq detects opposite deviations in different samples, the CNA is split up into two separate mutation, one for each affected allele, each tracked and assigned to a clone separately. An example of this is shown in **Supplementary Methods Figure 7**, where chromosome 8 is gained independently in the diagnosis and relapse sample of a donor with Acute Myeloid Leukemia (AML). Repeated mutations suggests convergent evolution and are important indicators of driver mutations. In this example, MYC is a strong tumor gene that is likely to confer an advantage to cell with trisomy 8.

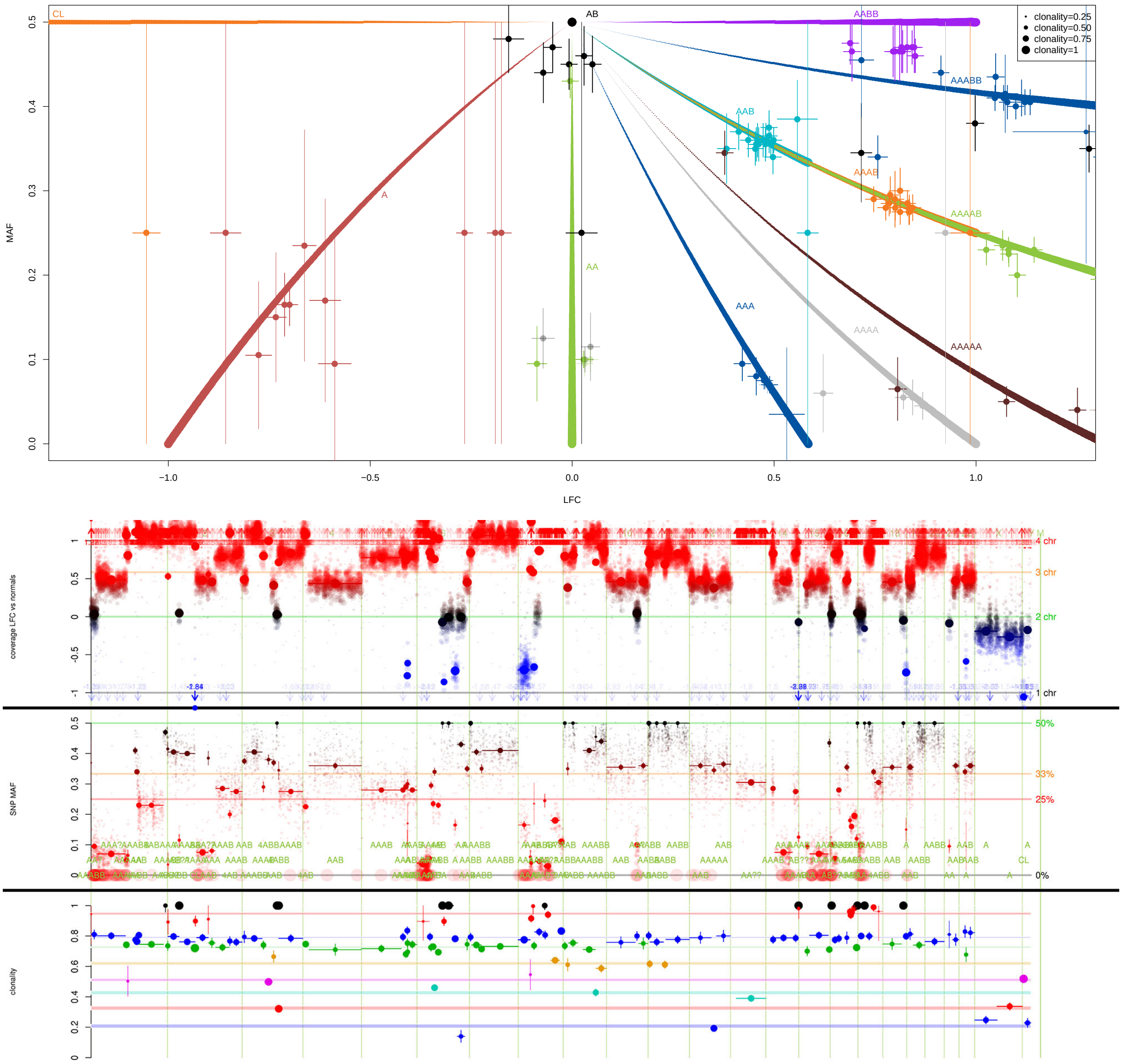


**Supplementary Methods Figure 6.** Illustration of the copy number and ploidy calling in superFreq. Top panel shows the log fold change (LFC) with respect to the reference normals on the x-axis and minor allele frequency (MAF) on the y-axis for each segment. Error bars show the propagated error for the LFC and MAF. Calling ploidy corresponds to shifting all data points along the x-axis. Trajectories show expected values for different copy number alterations at growing clonality. The colour of the data points match the trajectory of the copy number call. A large ploidy is called in this example, with most segment assigned to AAB (cyan), AAAB (orange) or AABB (purple) copy numbers. Bottom panel is CNA calls over the genome, showing LFC, MAF and clonality of the call. The size of the dots represent accuracy, based on the adjusted limma estimates for LFC, and based on the effective coverage for the BAFs. Segments, shown as dots with horizontal lines, also shows error estimates through an error bar and point size, and the extension of the segment on the x-axis. CNA calls are shown below the BAF segments, where uncertain calls (inconsistent data) are marked with "?" or "??". SuperFreq defaults to less exotic copy number states, so in this example an AAAB genotype at 100% clonality is selected rather than an AAAAB call with 80% clonality. The presence of other CNA calls around 80% clonality suggests that this call is incorrect. Data is from the Diffuse Large B-cell Lymphoma sample in donor P34 of [1].

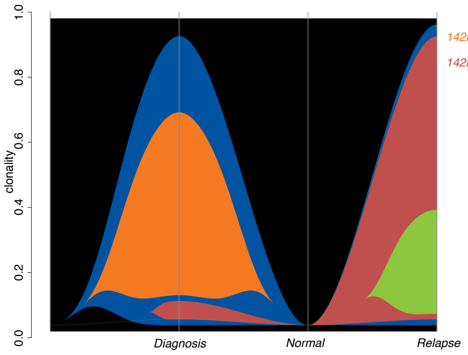


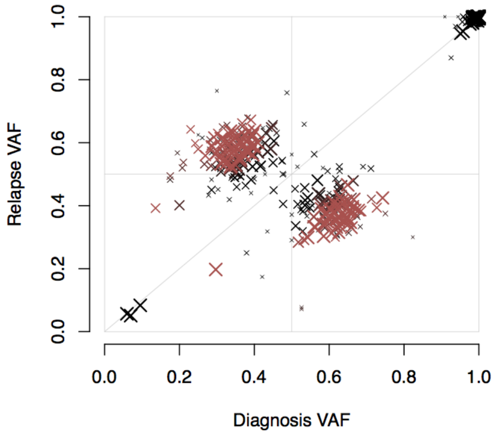


**Supplementary Methods Figure 7**. Left) River plot of AML.102[2]. The orange diagnosis specific clone and the red relapse specific clone (subclonally present in diagnosis) both have gain of chromosome 8, but with AAB and BBA genotypes respectively. Right) Scatter plot of variant allele frequencies of chromosome 8 population variants (ExAC allele frequency $> 0.01$) in AML.102. Variants with significantly different VAF (Fisher's exact test, Benjamini Hochberg multiple hypothesis corrected) in the diagnosis and relapse samples are coloured red. The off-diagonals clusters of heterozygous variants show that different alleles are affected by the CNA.

#### 1.7 Clonal tracking

##### 1.7.1 Clonality of somatic SNVs

The CNA calls are already made with a clonality and error estimate, but the SNV frequencies need to be converted to clonalities. In a normal diploid region, this is easy, as you just multiply the VAF by two, and estimate the error from a binomial distribution.

If a CNA is present over a SNV, there are three possible cases:

- **SNV subclone of CNA** The SNV can be present in a subset of the cells with the CNA. In this case, the SNV happened after the CNA, and will only be present on a single allele. There is an upper limit on the clonality (and thus VAF) of the SNV, as it cannot be larger than the clonality of the CNA.

- **CNA subclone of SNV** The CNA is present in a subset of the cells with the SNV. Here, the CNA happened after the SNV, and the clonality-VAF relation depends on which allele the SNV was on. In this case, there is a lower limit on the SNVs clonality, as it has to be as large as the CNV clonality.

- **CNA and SNV disjoint** The CNA and SNV are present in different cells. In this case, there is an upper limit to the clonality of the SNV, as the sum of the SNV and CNA clonalities cannot exceed 1.

These three scenarios are all tested, and if only one scenario is consistent with the clonality constraint, that option is selected. If multiple options are possible, the uncertainty of the SNVs clonality is increased to cover all possibilities. This inclusion of theoretical uncertainty can lead to SNVs of otherwise high quality getting large uncertainties, but we find this preferable over confident but occasionally incorrect clonality call. Often the variant is still assigned to the correct clone called from other mutations based on more certain clonality estimates in other samples from the individual.

##### 1.7.2 Clustering of mutations

**Anchor mutations** The most reliable mutations, SNVs and CNAs, are selected to anchor the clonal structure. Copy number calls smaller than 5Mbp and indels are excluded, as well as mutations with an average clonality error larger than 0.2. Somatic SNVs are required to have a somatic score $s$ larger than 0.95. These mutations are used for clustering and sets the phylogenetic structure. More uncertain mutations are afterwards associated to the clones determined by the anchor mutations. This allows superFreq to include error prone mutations and potential false calls, such as indels, low coverage SNV or CNAs over small regions, in the final output, with no risk of false clones being called from noise or miscalls.

**The germline clone** A germline clone is added to the analysis. This is an artificial mutation with clonality 1 and uncertainty 0 in every sample. The purpose of this mutation is to attract and identify germline mutations in the clustering, separating them from the somatic mutations.

**Clustering** The anchor mutations are clustered hierarchically. The distance measure $D$ between two sets $I, J$ of mutations is determined from how consistent the two sets are with an average clonality determined from the union $K = I \cup J$. The mean clonality $C_s$ over samples $s$ from all mutations $K$ is determined with the usual error-weighted mean in eq. 1.2. We first calculate a relative error $\sigma_{ks}$ for each mutation $k \in K$ and sample $s$:

$$\sigma_{ks} = |c_{ks} - C_s| / \Delta c_{ks}$$

where $c_{ks}$ and $\Delta c_{ks}$ are the clonality and clonality uncertainty of mutation $k$ in sample $s$. We use a normal distribution to convert the relative errors to probabilities, combine the probabilities with $I$ and $J$ separately with Fisher's method, and then convert these two probability back to relative errors $\sigma_{Is}$ and $\sigma_{Js}$ with normal distributions. The largest of these two relative errors across samples is used as distance metric between $I$ and $J$.

$$D_{IJ} = \max_s (\sigma_{Is}, \sigma_{Js}) .$$

Once no two clusters $IJ$ have $D_{IJ} < D_{\max}$ the clustering stops. $D_{\max}$ is a user adjustable parameter with default 2.3 corresponding to p $>$ 0.01 in a normal distribution. Each cluster is assigned a consensus clonality and uncertainty based on the anchor mutations in the cluster.

**Phylogeny** The clusters are interpreted as cellular populations and organised into a phylogeny. Sometimes multiple phylogenies fit the same clonality, and in those cases superFreq favours linear over parallel evolution. That is, superFreq prefers to group two clones as subclones rather than disjoint clones.

SuperFreq takes all clones and checks which can be subclones of which other clones. For clone $A$ to be a potential subclone of clone $B$, $A$ has to have a consensus clonality equal (with 1.5 uncertainty) or smaller than $B$ in all samples. This relationship is made antisymmetric (a pair of clones can't both be subclones of the other) by using the sum of clonalities to resolve ties. Then the relation is extended to its transitive closure (a subclone of a subclone is also a subclone).

Starting with the clone with the largest summed clonality, which will always be the germline clone, superFreq iteratively finds the phylogenetic tree. Each step identifies the subclones $A_s$ of a clone $A$ that can not be subclones of other subclones $A_s$, which are assigned as immediate disjoint subclones $A_d \subset A_s$. In order of summed clonality over all samples, superFreq iterates over $B \in A_d$, where all subclones $B_s$ of $B$ are assigned to $B$ and not used in other immediate disjoint subsets $A_d$. The enforced antisymmetry and transitivity of the subclonality relationship, together with the starting germline clone having all other clones as subclones, ensures that this algorithm always finish and includes all clones.

In this formalism, we expect unitarity: the summed clonality of all immediate disjoint subclones $A_d$ should be equal or smaller than the clonality of $A$ in every sample. If this is not the case (allowing for unitarity violations within errors), then all clusters of mutations corresponding to the clones cannot correspond to real cell populations. SuperFreq addresses this situation by determining which of the clones is most dodgy, removes that clone from the phylogenetic analysis, and starts over. Mutations in the dodgy cluster can still be assigned to other clones in the following section. The dodginess score $d$ is a sum of a range of warning signs

$$d = \frac{1}{N} + L + \frac{N_{\text{CNA}}}{N_{\text{CNA}} + N_{\text{SNV}} + 1} + B + U.$$

$N$ is the total number of mutations in the clone to tag clones based on a single or very few mutations. $L$ is the fraction of the distances between SNVs that are less than 10kbp, marking misalignment and localised contamination artifacts. $N_{\text{CNA}}$ and $N_{\text{SNV}}$ are the number of somatic CNAs and SNVs. Systematic errors in CNA calling, such as incorrect ploidy estimates or very noisy coverage profiles, can cause clones driven by a large number of CNAs but few or no SNVs. $B$ is -0.2 if the clone contains at least one CNA and at least one SNV, 0 otherwise. This gives a small discount to clones supported by both types of mutations. $U$ flags clones that are unchanged over samples $s$ by measuring the difference between the sample with the highest value and lowest value, allowing for uncertainty:

$$U = 2\left(0.5 - \left(\max_s(c_s - \Delta c_s) - \min_s(c_s + \Delta c_s)\right)\right).$$

A clone without any significant change will score $U = 1$, while a clone changing more than 0.5 in clonality will score 0. Negative values are increased to 0. The dodginess score encodes an automated quality control to identify the cluster of mutations that does not correspond to a cell population in case of unitarity violation.

This unitarity constraint can prove immensely powerful in removing clones driven by false calls. This is particularly useful when a matched normal is not present, where noise shows up at low to intermediate frequency mutations throughout all samples. These mutations are often grouped up into one or multiple roughly constant noise clones. Real clones in the samples that change from low to high clonality are not consistent with noise clones under the unitary constraint, and the noise clone can be removed through this process.

**Linking residual mutations** Once the clusters are finalised from the anchor mutations, lower quality mutations, SNVs, indels and CNAs, are compared to the existing clusters, and mutations that are consistent with a cluster within errors is assigned to the best fit. Mutations that do not match any cluster are discarded from the clonal analysis.

##### 1.7.3 Non-repeating mutation assumption and convergent evolution

Also known as the infinite site assumption, we assume that no somatic mutation happens twice, or is mutates back to reference. This assumption is only rarely broken, but those cases can be of great importance, potentially a sign of convergent evolution. There is no way to detect repeated mutations if the data is also consistent with a single mutations in an earlier clone. However, if there is no earlier clone that can consistently harbour the mutation, then the mutation will be filtered out in the clonal consistency step.

Considering the above, the signature of convergent evolution is a clone consisting of a single high confidence mutation (and potentially false calls attracted to the mutations) that is only present in the dodgy river output, but not in the default river that has undergone a clonal consistency filter. It should be possible to split the clone into two (or more!) clones that agree with existing clones in the default river output. This kind of analysis is capable of detecting driver mutations from a single cancer case, but should also be used with extreme caution.

#### 2 TCGA analysis

The data and code required to reproduce the figures and parts of the analysis are available at https://gitlab.wehi.edu.au/flensburg.c/superFreqPaper The available analysis is restricted by data sharing policy of TCGA germline information.

##### 2.1 Sample selection

We randomly selected 10 participants from each of the 33 projects in TCGA, requiring exactly one cancer and normal sample to facilitate unambiguous comparison to external calls on SNV or CNA calls. 26 participants were removed due to issues with meta data or file downloads. In all analysis superFreq was run with default parameters using 10 random normal TCGA samples with the same capture regions, not necessarily from the same tissue type. A list of the donors, including the reference normal samples, is available in participants.tsv on gitlab.

##### 2.2 Somatic SNV calls

The somatic SNV calls from SomaticSniper, MuSE, Varscan2 and MuTect2 were downloaded from GDC. For MuSE, Varscan2 and MuTect2, only variants with the quality filter PASS were used. No such filter was available for SomaticSniper, so all variant were used. SuperFreq SNVs with `somaticP` larger than 0.5 were used. All indels were removed as some of the methods only call SNVs.

**Supplementary Figure 1** shows the number of SNVs called by each combination of methods, divided by the number of consensus SNVs: SNVs called by all five methods. The plot is limited to 2 times the number of consensus SNVs. Participants with fewer than 10 consensus SNVs are excluded.

##### 2.3 Dilution and Slicing of the samples

We combined the cancer BAM file with the matched normal to create samples with lower cancer purity and different purities in different regions of the genome. First the average ploidy was calculated for the cancer and normal sample using the superFreq analysis, then the number of reads per ploidy was calculated for both samples. The sample with the larger value was downsampled so that both had the same number of reads per ploidy. This ensures that diploid regions are expected to have the same read depth in both samples, which allows for consistent mixing of samples.

The samples were then blended. A fraction $F$ of the reads was taken from the cancer sample and combined with a complementary fraction $1 - F$ of the reads from the normal samples into a single diluted BAM. F took values of 0.1, 0.3, 0.5, 0.7 and 0.9. Note that

F is not the purity of the blended sample, as the original cancer sample may not be 100% pure, and the normal sample may be contaminated.

The superFreq CNA calls larger than 10Mbp from the cancer-normal analysis were then sliced. A fraction of each of the genomic segments affected by copy number alterations is taken from the cancer sample, and the complementary region is taken from the normal sample. The size of the fraction is reduced by a factor 10 for each iteration, down to a minimum size of 100kbp. The segments are shrunk towards the captured gene closest to the center of the copy number segment, to ensure that at least one captured gene is inside the sliced copy number alteration.

##### 2.4 Comparison of Copy Number Calls

The three copy number callers: ASCAT, Sequenza and superFreq, were compared based on relative DNA abundance. The copy number call from ASCAT were lifted over from hg19 to hg38 using `segment_liftover`. First, we accounted for differences in ploidy estimate by dividing the DNA abundance with the called ploidy. Then each pair of overlapping segments was compared and classified as consistent if the relative DNA abundance was within errors. Error estimates of the $\log_2$ abundance from the two compared methods were added in square, and an additional error of 0.1 was added in square to account for uncertainty in the ploidy estimate. The fraction of genome in agreement is then calculated as the number of overlapping bases between agreeing segments divided by total number of overlapping bases. The error of the $\log_2$ DNA abundance was not provided in the ASCAT or Sequenza calls and was set to 0.1 for this purpose.

For the more stringent comparison we assessed allele sensitive copy number call and segmentation. For two overlapping segments to be classified as being in agreement, the size of the overlap had to be at least 80% of the size of the largest segment, and both methods had to call the same allelic copy number.

##### 2.5 SuperFreq Copy Number Sensitivity Assay

SuperFreq was run on the diluted samples and the sliced samples, and the copy number calls were compared to those obtained with the original cancer-normal pair. The calls were classified as detected if there was a copy number call over a segment with overlap to the true segment of 90% of the larger of the two segments.

##### 2.6 Clonal simulations

Only participants with at least one clone called by the superFreq cancer-normal analysis were included in the clonal simulations, which was 289 out of the 304 studied participants.

We devised a mixing approach to generate samples that could be used to test superfreq (outlined in methods, and schematically in**Figure 4a** in the main paper. There are some limitations to this approach, for example, chromosome 8 to 14 will be identical in Mix 1 and Mix 3 as it is just the normal samples, making it easier to track differences. On the other hand, the number of mutations in each clone is reduced, as the cancers somatic mutations are split across the four clones. This can pose a problem in particular for cancer types with low mutation burden, as there can be very few or no mutations to track in each clone. Further, any real subclones in the original cancer samples will propagate into proportional of the four constructed clones, making the system noisier. Cancer contamination in the normal sample will produce different clones that do not go all the way down to 0 clonality, and a low purity of the cancer sample will produce low clonalities of all clones in all samples. Taken together it is difficult to determine what clonalities should be expected as truth in the simulated samples. To account for these issues, we allow for subclones of the expected clones to be called without counting them as false positives, and we count any clone proportional to the expected clone as a match. That is, any detected clone present in sample one with clonality of 0 in the other two samples will count as match for the chr1-3 clone. The high accuracy of mutation assignment for both superFreq and SciClone suggests that this is a reasonable approach.

We allow the clonality to be within 2 error bars of expected values for superFreq, and correspondingly for SciClone we move the (asymmetric) upper and lower limits a factor 2 away from the estimated clonality. Both superFreq and SciClone were run with default settings. Default SciClone requires 10 variants (to consider up to 10 clones) with at least 100 read depth over normal diploid regions to run. When this requirement was not met, we decreased the minimum read depth to 30, and the maximum number of clusters to 5. In case that also fails, we run without copy number information, where all variants will be used independently of local copy number status. Cases where all attempts fail, such as samples with low coverage or low mutation rate, were excluded from the SciClone analysis. On the other hand, if superFreq does not find any mutations that pass the quality filters, then no cancer clones will be called and it is reported as a false negatives in the assay.
